# Phenotypic similarity is a measure of functional redundancy within homologous gene families

**DOI:** 10.1101/2022.07.25.501402

**Authors:** Jessica A. Comstock, Merrill E. Asp, Fatmagül Bahar, Isabella Lee, Alison E. Patteson, Roy D. Welch

## Abstract

Robustness to the impact of mutation can mitigate phenotypes that have the potential to inform gene function. This robustness is often encoded into the genome through gene duplication, among other mechanisms. Duplication is a source of structurally similar genes that can retain some functional overlap as they diverge, and as such contribute to functional redundancy in the face of mutation. While redundancies have been explored in groups of two or three paralogs by generating double and triple mutants, it is unclear to what extent larger homologous gene families contribute to robustness through functional redundancy. Here, we used phenotypic similarity as an indicator of functional redundancy to explore the extent to which homologous gene families contribute to redundancy in function. We hypothesize that, since functional redundancy is more likely to occur within gene families where genes are structurally similar, mutant strains within the same gene families would be more phenotypically similar. We generated 265 single-gene disruptions in four homologous gene families of *Myxococcus xanthus*, used time-lapse microscopy to generate time series of multicellular development, and developed an image analysis pipeline to compare phenotypic characteristics among different strains. We show that mutant strains cluster by gene family in the phenotypic feature space with principal component analysis, demonstrating that families of homologs can contain extensive functional redundancy networks.

## Introduction

A reverse genetics approach to characterizing a gene often begins by disrupting or deleting the gene and observing the resulting phenotype. Differences between the mutant and wild-type phenotypes can provide invaluable insights regarding gene function(s), but in practice many single-gene knockouts, even those in genes predicted to be important based on previously studied homologs, yield phenotypes that are relatively minor or indistinguishable from the wild-type organism (Diss *et al*, 2014; Giaever *et al*, 2002). This robustness to the phenotypic impact of genetic mutation is an important part of an organism’s phenotype and has implications for fitness.

Robustness is commonly attributed, at least in part, to functional redundancy, or the tendency for functionally similar genes to compensate for the role of a disrupted gene (Ohno, 1970). Functional redundancy can arise through many mechanisms including duplication and divergence, where reduced selective pressure can cause paralogs to accumulate mutations and take on new, slightly different functions over time (Vandersluis *et al*, 2010; Krakauer & Plotkin, 2002). Paralogs that are maintained over long timescales often retain some of their ancestral function in addition to their diverged function (Kuzmin *et al*, 2022; Dean *et al*, 2008), thus building in redundancy. Repeated gene duplication events can give rise to large gene families wherein genes have a range of biochemically similar but specialized functions. Though many homologs in a gene family may be capable of performing a similar function, due to divergence it is difficult to predict which genes might be able to compensate for the function of others. The most recent duplicates within a gene family are not always capable of being functionally redundant while some older and more diverged paralogs are (Baker *et al*, 2021). Sequence similarity alone is not enough to predict functional redundancy, and the extent to which duplicates contribute to robustness varies across organisms (Hannay *et al*, 2008). For these reasons, it is unclear to what extent families of homologs are contributing to the functional redundancy that gives rise to robustness in biological systems.

Many studies attempting to elucidate functional redundancy in the genome involve the creation of single and double knockouts of paralogs to probe for synthetic lethality (Thomaides *et al*, 2007; diCenzo & Finan, 2015; Butland *et al*, 2008). While this method is effective in assessing functional redundancy in pairs of closely related genes, it is limited in its power to explore larger networks of redundancy, as may exist in expanded gene families. A double mutant that does not show a more significant phenotype than each of the corresponding single mutants could imply either that the genes are non-redundant, or that they are part of a larger redundancy network that has a strong buffering capacity and therefore has decreased fragility in the face of genetic perturbation (Lehár *et al*, 2008). In this way, phenotype is often the readout for assessing redundancy and robustness within biological systems. The phenotypic impact of mutation reveals information about robustness, and we can investigate the mechanisms that lead to robustness by considering gene sequence, so understanding how redundancy affects robustness is a crucial genotype-to-phenotype question.

Any given gene processes the flow of information from precursors, producing outputs that feed into other networks or cellular functions. In a simple case of non-redundancy, a gene produces one protein with a primary function, and when this gene is intact, expresses a wild-type phenotype (Fig. 1A). A mutation in this gene would severely impact the fitness of the organism. However, if a given gene is part of a network of structurally similar genes which each have their own primary function but also retain some ancestral function, as in gene families that arise from duplication, the impact of a mutation can be diffused through the other members of its network, producing a relatively minor deviation in phenotype. Redundancy networks (Fig. 1B), which we here define as the group of two or more genes whose products can compensate for the loss of function of one another, allow for the rerouting of information through alternative pathways so that the end result has a minimal impact on fitness. As shown in Figure 1C, an additional byproduct of this buffering effect is that knocking out one member of a redundancy network should produce a similar phenotype as knocking out any other member of that group, because the entire set of genes is affected no matter which component of the network is disrupted by mutation. In this way, phenotypic similarity may be an important indicator of functional redundancy within homologous gene families and may provide insights into the level of robustness in a genome. Further, because a gene’s redundancy network likely overlaps significantly with its family of homologs due to the relationship between protein structure and function (Fig. 1D), we predict that mutations in genes from within the same gene family will be more phenotypically similar.

**Figure 1.**
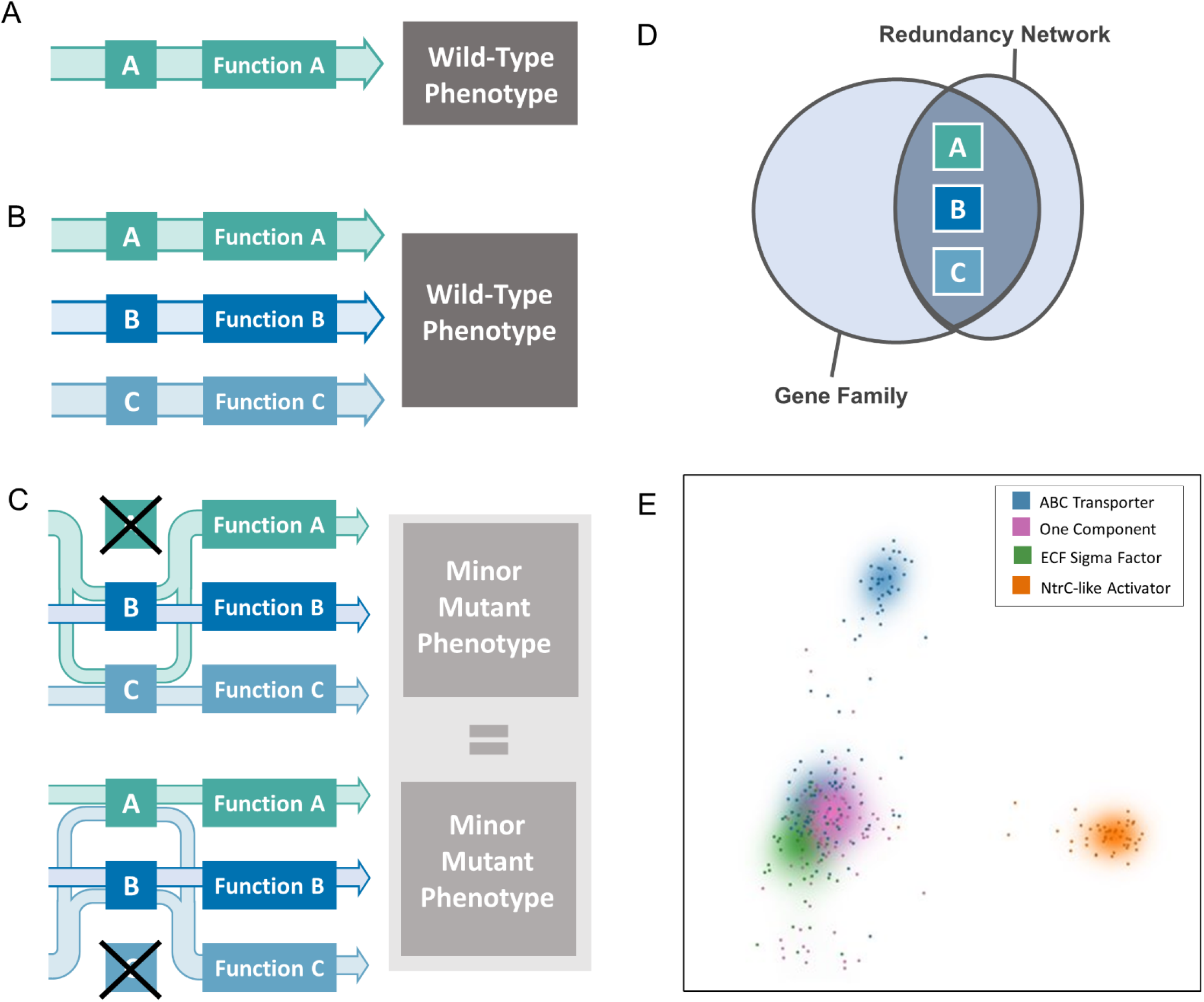
Functional redundancy resulting in phenotypic similarity. **(A)** In a pathway with no redundancy, Gene A contributes to Function A. Any mutation that renders Gene A nonfunctional would produce a severe phenotype or lethality if Function A is essential. **(B)** Genes A, B, and C belong to the same redundancy network, meaning each gene can compensate for the loss of function of one member of its network. In the scenario where all three genes are functional and operating optimally, each gene contributes to its primary function (for example, Gene A is responsible for most of the contribution to Function A), producing the wild-type phenotype. **(C)** When a mutation occurs that renders Gene A nonfunctional (top), the input to Gene A gets rerouted through Genes B and C such that Function A can still occur, but in a slightly reduced capacity (indicated by thickness of arrows compared to panel B). Since Genes B and C are processing more input from A, Functions B and C are also affected and operate at a reduced capacity. The slight reduction in function of all three network components produces a phenotype that is relatively minor and may be indistinguishable from wild-type. A mutation in Gene C (bottom) would result in a similar phenomenon, where the input that normally feeds into Gene C is processed by Genes A and B, resulting in overall decreased output from each. In this model, a mutation in one member of a redundancy network affects the output from all components regardless of which gene contains the mutation (indicated by the similar output arrow size of top and bottom of panel C), and we predict that mutations in members of the same redundancy network will produce similar phenotypes. **(D)** Though not every member of a gene family is functionally redundant, and there may be redundant genes that do not belong to the same gene family, the relationship between structure and function of proteins dictates that genes in the same redundancy network are likely to come from the same gene family. If redundancy networks come primarily from the same gene family, and components of redundancy networks show similar mutant phenotypes, then members of the same gene family would be more likely to produce the same mutant phenotype. **(E)** Similarity of the protein sequences used in this study by multidimensional scaling. Each point represents one gene, with a Gaussian kernel density estimate to guide the eye. Proteins that are more similar in sequence, belonging to the same gene family, cluster together. Each gene family forms a single cluster with the exception of the ABC Transporters, which form two major clusters due to the different subunits. The highly conserved ATP-binding domains (Rees *et al*, 2009) separate very distinctly from the periplasmic and substrate-binding domains. We predict that mutations in genes that belong to the same gene family will be more phenotypically similar than they will be to mutant phenotypes in other paralogous gene families. Thus, we expect phenotype to cluster by gene family.

To test this, we phenotypically characterized over 250 single-gene mutations in *Myxococcus xanthus*, a soil bacterium with a large genome containing multiple homologous gene families (Goldman *et al*, 2006) and examined the relationship between gene family and phenotype. Under nutrient stress, swarming cells of *M. xanthus* undergo development, aggregating into multicellular fruiting bodies wherein populations of cells will differentiate into spores (Bretl & Kirby, 2016) (Fig. 2A). Since the ability of *M. xanthus* to form fruiting bodies and sporulate is directly tied to its fitness, it is likely a robust biological process that involves many functionally redundant genes. We generated a library of microscopic time-lapse movies (time series) showing the development of 265 knockout strains of *M. xanthus* belonging to four different gene families (102 ABC transporter genes, 45 NtrC-like activators, 80 One component signal transduction genes, and 38 ECF sigma factors; see references Yan *et al*, 2014, Caberoy *et al*, 2003, and Abellón-Ruiz *et al*, 2014 for previous work on some of these genes in *M. xanthus)*. We made qualitative observations of the ways in which resulting phenotypes differed from wild-type and used these observations to inform a novel image processing and phenotypic analysis pipeline that automates quantitative measurements of phenotype that are explicitly defined. Although previous studies have used image processing to extract phenotypic features of aggregate formation (Xie *et al*, 2011), this work has applied these tools to the largest library of time series of which we are aware, necessitating a new pipeline and analysis methods. Finally, we compared the similarity of phenotypes across gene families using principal component analysis (PCA). Wefound that, just as mutant strains within a gene family cluster by sequence similarity through multidimensional scaling (Fig. 1E), they also cluster by gene family in the phenotypic feature space with a statistically significant sharpness (i.e. small cluster size) and separation of clusters, indicating large networks of redundancy within these gene families.

**Figure 2:**
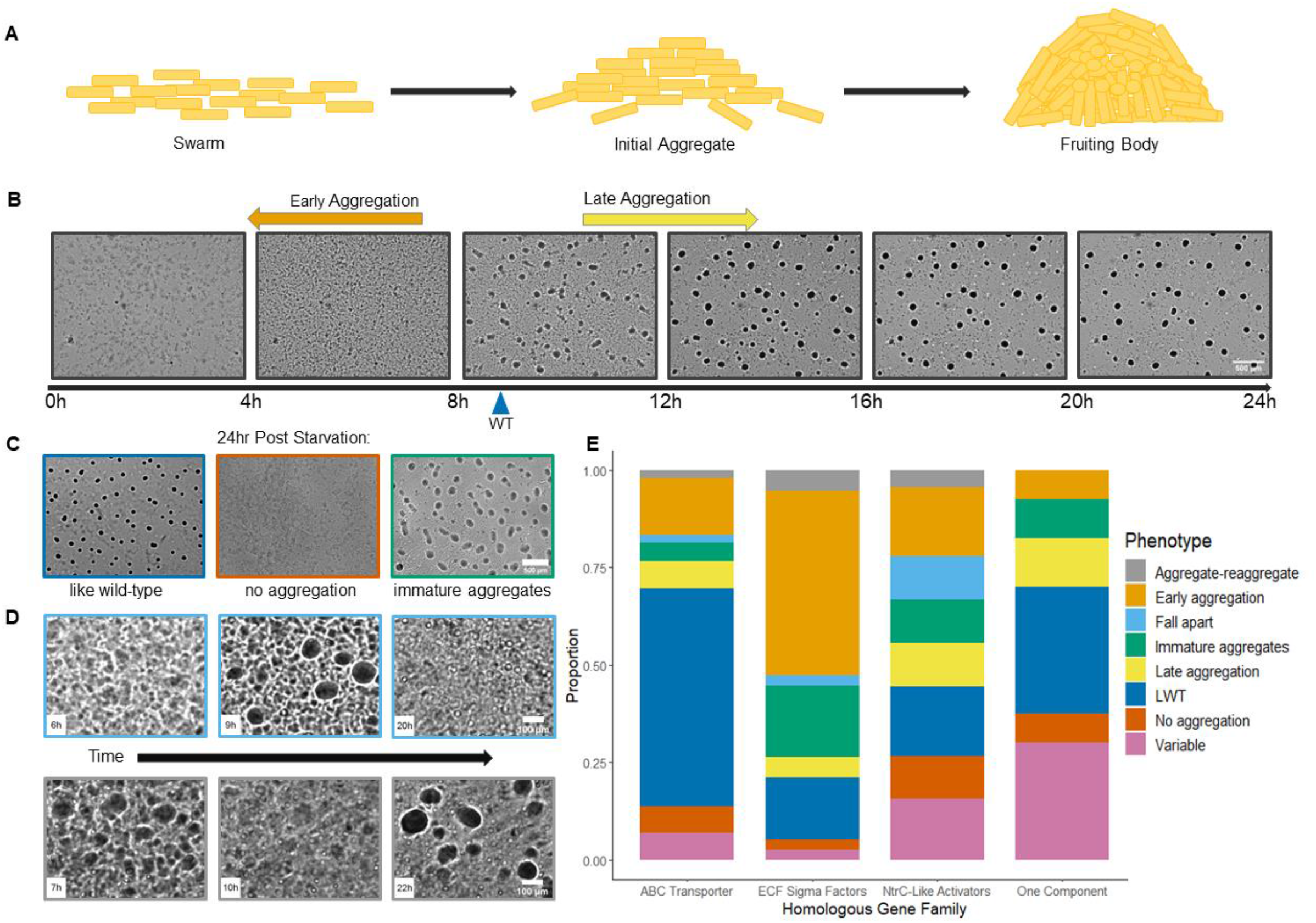
Manual categorization of *M. xanthus* development. **(A)** Upon sensing nutrient stress, vegetative *M. xanthus* cells undergo a developmental process that culminates in spore-filled fruiting bodies. **(B)** Wild-type *M. xanthus* cells on TPM agar begin to cluster into early aggregates after 9 hrs of starvation (blue arrow), and as more cells join the premature aggregates over the course of 24 hours, the aggregates mature into fruiting bodies that appear round and dark with conventional brightfield microscopy. Mutant strains that show initial aggregation either before or after the average time for wild-type are assigned the early aggregation (orange arrow) and late aggregation (yellow arrow) phenotypes, respectively. Scale bar 500 µm. **(C)** Like wild-type (LWT) mutants that produced dark, circular fruiting bodies on a timeline similar to wild-type (left), non-aggregating mutants (center), and mutants that produced immature aggregates (right). Scale bar 500 µm. **(D)** Some mutants formed initial aggregates that eventually shrank and fell apart (top). Other mutants formed initial aggregates that fell apart before re-aggregating into mature fruiting bodies (bottom). Scale bar 100 µm. **(E)** Distribution of development classifications within each gene family.

## Results

### Manual Characterization of Development Phenotypes

Under starvation, a swarm of *M. xanthus* cells will execute a developmental program during which millions of rod-shaped cells coordinate their movements and self-organize into dome-shaped multicellular aggregates. Some nascent aggregates destabilize and disperse, but most persist and continue to grow; when the persisting aggregates become large enough, cells in the middle of each differentiate to form a cluster of spores, at which point they are considered mature fruiting bodies (Fig. 2A). Capturing this process with time-lapse brightfield microscopy results in a time series of grayscale images where initial aggregates appear roughly circular with irregular boundaries, somewhat darker than the background swarm. Later in the time series, dispersing aggregates shrink and disappear, and the persisting ones grow and darken, with boundaries that become stable and clearly defined (Fig. 2B). Image features such as these can be leveraged to compare development phenotypes between wild-type and mutant *M. xanthus* strains.

For this study we recorded 24-hour time series for wild-type and a set of 265 single gene knockout mutant strains (Table S3), with an average of three replicates per strain. Due to their important roles in signal transduction, transport, and transcriptional regulation, we predict that genes within these families will be part of redundancy networks to ensure robustness. We compared the mutant phenotypes to our wild-type strain with an emphasis on aggregate composition and dynamics. Wild-type aggregation initiated at 9.2±1.6 hours and formed uniformly dark circular aggregates with stable and clearly delineated boundaries within 24 hours (Fig. 2B). Mutant strains that consistently initiated aggregation either before or after wild-type were designated “early” or “late”, respectively. Mutant strains that consistently initiated aggregation at the same time as wild-type but had aggregates that failed to darken and/or form clearly delineated boundaries were designated “immature” (Fig. 2C). Mutant strains that initiated aggregation at the same time as wild-type but then all the aggregates dispersed within 24 hours were designated “fall apart” (Fig. 2D). Mutant strains that initiated aggregation at the same time as wild-type but then all the aggregates dispersed and then re-aggregated within 24 hours were designated “aggregate-reaggregate” (Fig. 2D). Mutant strains that consistently matched all aggregation criteria and were indistinguishable from wild-type were designated “Like Wild-Type” (LWT). Finally, Mutant strains where the replicates displayed different developmental classifications were designated “variable”.

### Distribution of Manual Development Phenotypes within each Gene Family

Of the mutant strains characterized in this study, less than 10% failed to initiate aggregation at all, and 62% consistently produced fruiting bodies that were qualitatively comparable to wild-type by the end of the 24-hour window. An additional 20% of mutants were able to initiate aggregation, but aggregates remained immature; some of these strains may have formed mature aggregates if the time series extended longer than 24 hours.

We hypothesize that the relatively high success rate of aggregation in these mutants is due, at least in part, to *M. xanthus* development being a robust phenotype. If redundancy networks are contributing to functional redundancy to produce this robustness, then, according to the hypothesis portrayed in Fig. 1, mutants within the same gene family will be more phenotypically similar. As an initial test of our hypotheses, we sorted the mutant strains into their gene families and visualized the proportional representation of our developmental phenotype classifications (Fig. 2E). The distribution of some phenotypes did seem to favor specific gene families. For example, LWT strains made up over half of the ABC Transporter family, the early aggregating strains compose nearly half of the ECF sigma factor family, and about one third of the One component family produce variable phenotypes in different replicates.

The manual categorization of development phenotypes presented here serves two purposes. First, it provides support for our hypothesis that *M. xanthus* development is as robust a phenotype as we expected, making it a suitable phenotype for observing the extent of functional redundancy networks in gene families. Second, though we do not claim these data alone provide sufficient evidence for the existence of redundancy networks, as the data show only the most obvious associations between gene family and phenotype, these qualitative observations do provide information about the various ways in which phenotype can differ during development. This was used to inform a more systematic, quantitative, and multidimensional characterization pipeline to test our remaining hypothesis about phenotypic similarity among families of paralogs: if redundancy networks contribute to robustness, and if those networks are comprised primarily of genes within the same family, then a grouping of mutant strains based on phenotypic features should also group the strains according to gene family.

### Automated Characterization of Phenotypes

We developed and implemented an automated image processing pipeline in Python (see Methods, SI). Using it, we were able to identify and track every aggregate in all the time series, recording changes in aggregate number, position, size, shape, and gray value. In total, our pipeline captured the developmental dynamics of more than 150,000 aggregates, both dispersing and persisting. These data were analyzed to determine the timing and position of significant changes in swarm dynamics, such as the initial onset of aggregation, the average aggregate growth rate, and the rate of change in aggregate gray value; these quantitative features serve as an unbiased and more accurate replacement for the manual phenotypes “early”, “late”, “immature”, “LWT”, and “variable”.

We identified 18 quantitative features (Fig. 3) to represent and measure the variation observed in the wild-type and mutant strains. For each time series, we calculated a list of these 18 numbers, mapping it to a single point in an 18-dimensional feature space. Distance between points in this feature space is a measure of phenotypic dissimilarity. To reduce the complexity of this data and visualize it, we used principal component analysis (PCA), a deterministic method with no additional input parameters, to reduce the feature space to two dimensions, PC1 and PC2. The distribution of points on a two-dimensional map of PC1 versus PC2 captures the phenotypic features that vary the most across the dataset, while discarding those combinations of features that vary less.

**Figure 3:**
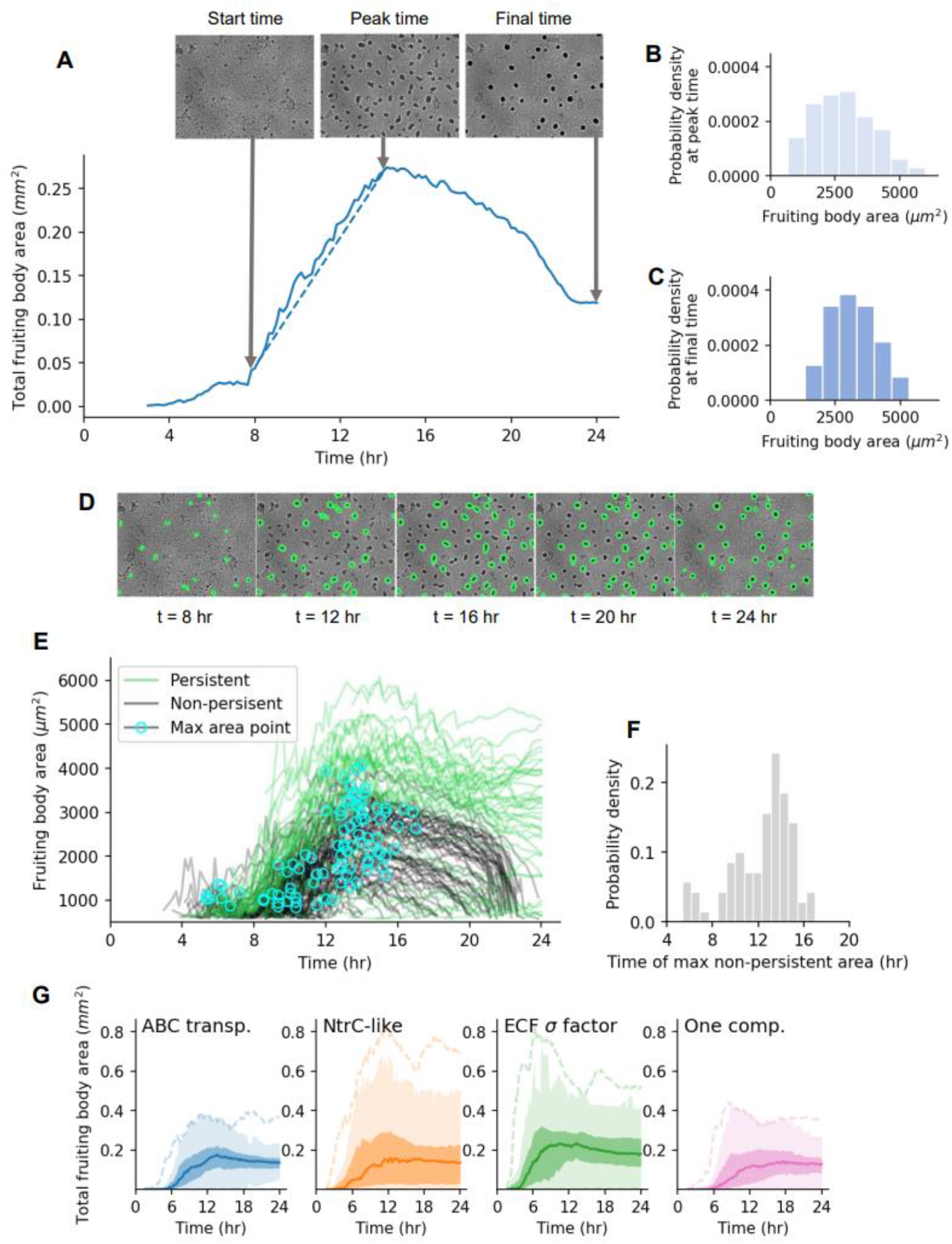
Automated quantification of fruiting body formation phenotypes. **(A-C)** Features related to global fruiting body development **(D-F)** Features related to fruiting body fate **(A)** A representative curve showing total fruiting body area over time in a 7.2 mm² field size. Images are shown of aggregation at start time, peak time, and final time (24 hours), all measured as time elapsed since inoculation (t=0). The slope of the dotted line in **(A)** gives the average growth rate, a key phenotypic feature. **(B)** Representative histograms from the same time series of average fruiting body area at peak time and **(C)** final time. The mean and standard deviation of these distributions are key phenotypic features. **(D)** Five representative time lapse images show fruiting body fate, either to persist or disappear after 24 hours of development. **(E)** Area versus time curves for each identifiable fruiting body in the same time series. For non-persistent fruiting bodies, the point of peak area is marked with a cyan circle. Two key features are the fraction of total identifiable fruiting bodies that persist (in this case, 32%, or 42 of 132), and **(F)** the standard deviation of the time at which non-persistent fruiting bodies peak in size (temporal coherence). Developmental dynamics can distinguish between time series of different homologous groups, as illustrated in **(G)** the curves for median total fruiting body area over time (quartiles bound the shaded regions, and outliers are bounded by the dotted lines). These variations are captured by 18 phenotypic features, with quantitative definitions given in Table S2.

The most significant phenotypic features are revealed by the makeup of the first two principal components, PC1 and PC2 (Table 1). These two components together account for 43% of the total variance. The constituent parts of both principal components represent a broad array of many different features, with no single outstanding feature. However, there are significant differences between PC1 and PC2. PC1 primarily represents growth rate, mean and standard deviation in fruiting body area at peak time, and mean and standard deviation in fruiting body area at final time. PC2, while sharing mean and standard deviation in area at peak time with PC1, also represents features involved with timing, including growth time, peak time, and temporal coherence. These developmental features with definitions are illustrated in Fig. 3. Although PC2 shares some key features with PC1, its correlations are different. For example, a high value in PC2 indicates high standard deviation in aggregate area and low growth rate, whereas a high value in PC1 indicates high standard deviation in fruiting body area and high growth rate.

**Table 1:** Makeup of PC1 and PC2 by phenotypic feature. Each primary component is a direction or vector in the 18-dimensional phenotype space, with its makeup shared to varying degree by each feature, with either a positive (blue) or negative (red) correlation. PC1 captures the direction of greatest variance in the overall dataset, and PC2 is the direction perpendicular to PC1 that captures the next greatest amount of variance. The features most strongly represented in each primary component are those that have the greatest potential to distinguish time series phenotypically across the dataset. Each feature is numbered according to its prevalence in PC1.

| PC1 Feature Name | Value | PC2 Feature Name | Value |
| --- | --- | --- | --- |
| 01) Final area mean | 0.363 | 07) Peak time | 0.426 |
| 02) Final area std | 0.325 | 05) Peak area std | 0.375 |
| 03) Growth rate | 0.322 | 13) Growth time | 0.328 |
| 04) Peak area mean | 0.321 | 04) Peak area mean | 0.322 |
| 05) Peak area std | 0.297 | 16) Temporal coherence | 0.305 |
| 06) Maturation rate | 0.279 | 02) Final area std | 0.298 |
| 07) Peak time | -0.256 | 17) Stability time | 0.229 |
| 08) Gray value % change | 0.254 | 03) Growth rate | -0.195 |
| 09) Max N falloff | -0.221 | 15) Start time | 0.193 |
| 10) Fraction of evaporators | 0.217 | 11) Lifetime std | 0.186 |
| 11) Lifetime std | 0.209 | 01) Final area mean | 0.172 |
| 12) Max N | 0.205 | 06) Gray value rate | -0.165 |
| 13) Growth time | -0.205 | 18) Lifetime mean | -0.141 |
| 14) Max mean area falloff | -0.118 | 08) Gray value % change | -0.133 |
| 15) Start time | -0.106 | 10) Fraction of evaporators | -0.132 |
| 16) Temporal coherence | -0.077 | 09) Max N falloff | 0.086 |
| 17) Stability time | -0.062 | 14) Max mean area falloff | -0.058 |
| 18) Lifetime mean | -0.045 | 12) Max N | -0.007 |

Each of the four homologous gene families used in this study has a different number of genes, and therefore each family is not represented by an equal number of mutant strains. This presents a potential bias towards over-represented gene families if PCA were to be performed on the entire dataset. To address this, we performed the PCA multiple times on random samplings of the time series such that each gene family is always equally represented (see Methods). We found that across all such samplings of the full dataset of over 1000 time series, the gene families always form clusters in distinct parts of two-dimensional phenotype space (Fig. 4), with clusters representing a different “typical” developmental phenotype for each gene family. There was significant overlap between the clusters, so that the differences between clusters only became apparent when using a sufficiently large sample size to visualize an estimate of the probability density. The PCA analysis therefore agrees with the manually derived developmental classifications presented in Fig. 2, in that mutant strains from each gene family display a full spectrum of phenotypes, but there are specific phenotypes that each family exhibits with higher frequency.

**Figure 4:**
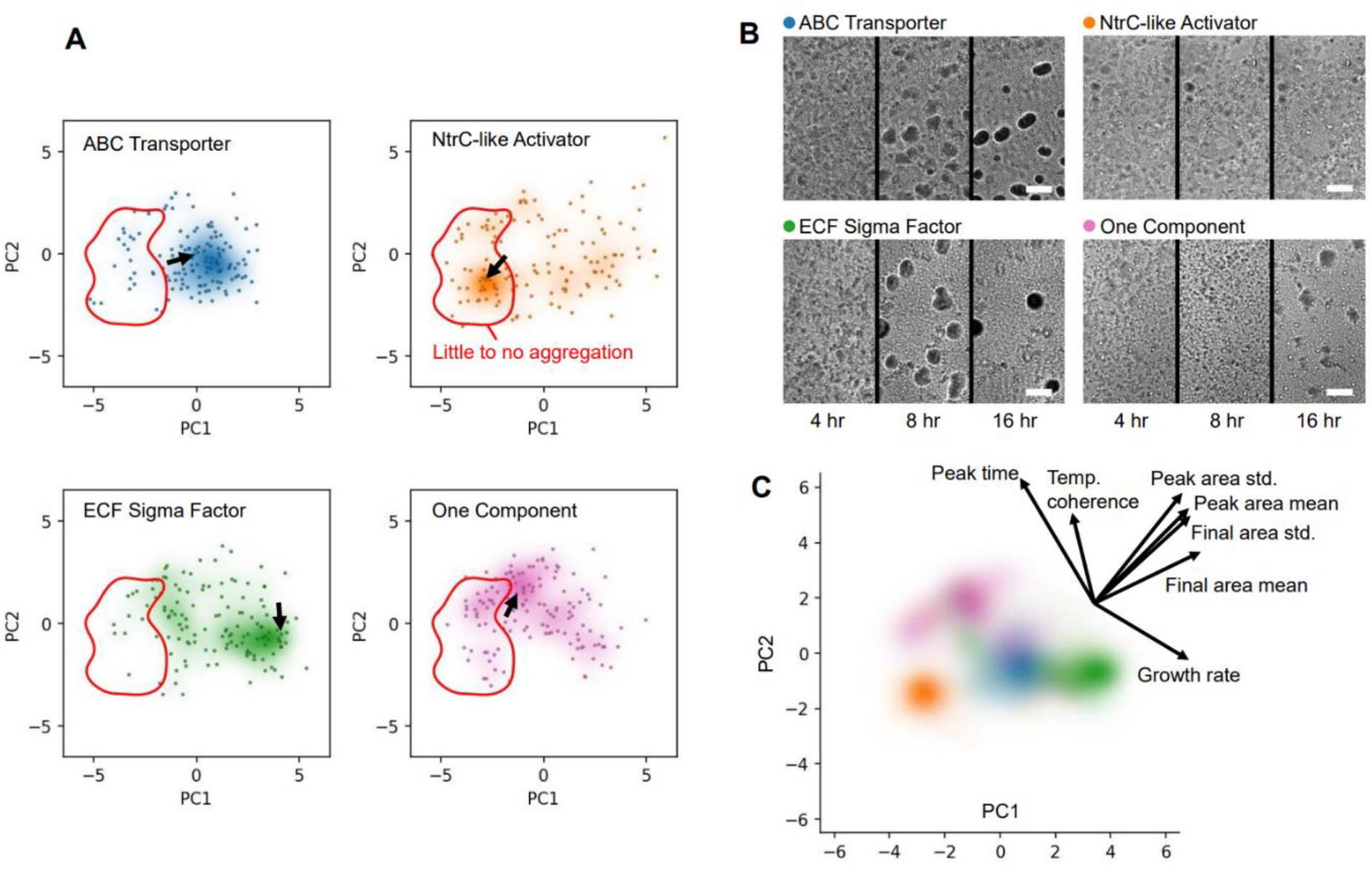
PCA reveals typical phenotypic features for each homologous family. **(A)** Each point represents a single time series, placed by phenotype according to values of PC1 and PC2. Units for PC1 and PC2 are arbitrary, but (0,0) represents average behavior. Behind points is displayed an estimation of the probability distribution function, using Gaussian kernel density estimation. Higher probability is plotted with higher opacity, revealing phenotypic clusters in each gene family. Outlined in red is a phenotypic zone containing only time series that exhibit little to no aggregation, a severe phenotype. Outside the red zone are successful fruiting body formation time series. An arrow points to a time series typical of the cluster, shown in: **(B)** The typical phenotype in each gene family cluster, illustrated with three frames from a representative time series, taken at 4 hours, 8 hours, and 16 hours after inoculation. Scale bars 100 μm. **(C)** Only the probability distribution estimates for each gene family are shown, illustrating both separation and overlap in phenotypic behavior. The directions of seven key phenotypic features are shown to indicate the coupled meaning of PC1 and PC2. Values of each feature increase in the direction of each respective arrow, with the length of the arrow indicating how much motion in feature space is caused by a fixed increase in the value of that feature, i.e. how significantly the feature is expressed by the two principal components.

The phenotypic cluster of ABC transporter mutants and ECF sigma factor mutants each show a distinct but consistently successful method of fruiting body formation, whereas One component mutants tend to vary widely phenotypically or form immature aggregates, and a plurality of NtrC-like mutants fail to form fruiting bodies entirely, as shown in the insets in Fig. 4B. Expanding on the comparison between the automated and manual phenotyping, we see that the phenotypic PCA clusters are not only consistent with the results of Fig. 2, but may also explain why a particular phenotype, such as “early aggregation” for the E sigma factor mutants, is expressed more often than for other homologous groups: a set of redundant genes robustly produces a minor mutant phenotype which shares multiple features in their aggregation formation dynamics, among which is an early aggregation time. Variability in phenotype can now also be measured by the spread of replicates in phenotype space, instead of needing to be classed as a separate phenotype unto itself. Representative visualizations and a basic measure of the spread of strain replicates are presented in Fig. S2.

There are two generic features of the way in which phenotype clusters form for each gene family. First, the separation between clusters indicates that mutations in each homologous group affect phenotype in a distinct way. Second, the size of each cluster’s individual peak points to how often each similar phenotype is expressed within the homologous group. Clusters in the PCA output were shown to be both separated and small with high statistical significance (with p-values of 10^−5^ and 0.0083 respectively) when compared to random groupings of time series instead of grouping by homologous group (Figure S1).

By manually reviewing each time series that fell near a phenotypic cluster, we determined that each represented a coherent overall phenotype with only a small number of outliers. Significantly, a developmental phenotype that could be considered severe, little to no aggregate formation, was shown to vary throughout the dataset, and the NtrC-like activators’ typical phenotype was shown to be a distinct form of failure to aggregate in which there was still bacterial motility, but little to no departure from a uniform swarm layer. Details on each typical phenotype and the metric values that distinguish them are available in the SI.

Our decision to characterize developmental phenotype via aggregate dynamics may have had a negative impact on our ability to differentiate the severe phenotypes that exhibit little to no aggregation. There is an identifiable phenotypic zone in the PCA output that represents little to no aggregation, as outlined in red in Fig. 4A. However, even within this zone, different behaviors are still distinguishable, and there is separation between homologous gene families. This is shown by the NtrC-like activators and One component mutants tending to occupy different phenotypic territory within the red outline. Differences in the dynamics of small, transient aggregates still hold significance in identifying typical behaviors across gene families.

Our observation that PCA of phenotypic metrics recapitulates homologous families of mutant strains indicates that phenotypic similarity and sequence identity are positively correlated across a genome, but this correlation does not scale down to fine-grained genetic differences. We observe that, within a homologous gene family, pairwise sequence similarity in this dataset correlates only weakly with phenotypic similarity, and therefore it is not an effective predictor of phenotype between pairs of genes within the same family (Fig. S3), consistent with previous findings (Hannay *et al*, 2008). The phenotype clusters for each gene family are populated by replicates of many mutant strains in that family, both the genetically similar and dissimilar. For the ABC transporters, 56% of strains (20 of 36) have replicates within the phenotypic cluster. This is also true for 24% (8 of 34) of NtrC-like activator strains, 39% (15 of 38) of ECF sigma factor strains, and 64% (16 of 25) of One component strains. Fine-grained genetic similarity is not necessary for redundancy to exist, and redundant gene networks may be quite large.

Taken together, Fig. 4 demonstrates that different homologous gene groups are likely to contain redundant gene networks, which each produce a distinct minor mutant phenotype.

## Discussion

Goldman et al proposed that the expansion of the *M. xanthus* genome due primarily to duplication and divergence has led to an enrichment of some gene families, especially those involved in cell signaling and transcriptional regulation, over others (Goldman *et al*, 2006). This asymmetry of enrichment is notable because it implies a purpose for the expansion of specific gene families. We propose at least part of that purpose is to create functional redundancy networks that act as buffers to stabilize *M. xanthus* development (i.e. robustness). In this study we confirm that *M. xanthus* development meets the criteria of a robust phenotype, by showing that more than 250 mutant strains with disruptions in genes that are part of four large homologous families display severe developmental phenotypes very infrequently. We then provide support for our hypotheses regarding the existence of redundancy networks by quantifying the phenotypes of these mutant strains and observing that using PCA to map phenotypic feature space also clusters the mutant strains according to the four homologous gene families.

Paralogs within a gene family may share similar molecular mechanisms, but they are expected to have different biological functions. For example, the ABC transporters in *M. xanthus* all perform active transport across membranes, but they are expected to transport different substrates. This would mean that a disruption of any one ABC transporter would cause a change in developmental phenotype specific to its substrate. There is no obvious reason why mutant strains of the ABC transporters would display similar changes in phenotype unless there is significant functional redundancy between transporters. There is also no obvious reason why the phenotypic similarities would include a plurality of a large homologous gene family unless the functional redundancy is widely distributed.

When a group of functionally redundant genes absorbs the effect of one member’s disruption with low overall stress on the system, the impact on phenotype is more subtle. The ABC Transporter and ECF sigma factor gene families exemplify this, as there are very few single gene knockouts that result in severe phenotypes (Fig. 2E). The phenotypes of those genes cluster sharply by homologous family in the PCA feature space (Fig. 4A), meaning that both gene families display a typical phenotype that is different from the others. Plausible biological explanations for this type of widely distributed functional redundancy can be made. Many ABC Transporters, due to varying homology in periplasmic and substrate-binding domains across the gene family, may be able to transport similar and/or overlapping substrates (Orelle *et al*, 2019; Durmort & Brown, 2015), mitigating the effect of many of the mutations in this gene family and most often producing like wild-type phenotypes. Similarly, robustness has been shown to be encoded in transcriptional regulatory networks by alternative pathways (Wagner & Wright, 2007), and though some studies suggest that alternative sigma factors display minimal crosstalk (Rhodius *et al*, 2013), it is not unprecedented for there to be overlap in the regulation of genes by multiple ECF sigma factors, creating networks of integrated regulation (Mascher *et al*, 2007; Luo & Helmann, 2009; Paget, 2015). Our data indicate that *M. xanthus* may use such networks of crosstalk among ECF sigma factors to coordinate transcription in response to extracellular signals, and that this may involve integration from many redundant or parallel pathways, ultimately leading to earlier aggregation initiation time and faster fruiting body growth rate than we see in wild-type for the majority of ECF sigma factor mutants.

In contrast, NtrC-like activator mutants show more severe phenotypes and cluster in a region where strains do not form fruiting bodies (Fig. 4). Though it could be argued that a non-aggregating strain indicates a lack of redundancy for the mutated gene, this seems unlikely given that the NtrC-like activators fail to produce aggregates in a way that is distinct from non-aggregating mutants in other gene families (Fig. 4A—region outlined in red). This again points to the idea of networks of redundancy, but highlights that there can be a cost to redundancy in some situations. Extensive research has shown that kinase-response regulator pairs tend to be very insulated with limited crosstalk, and that this feature rapidly evolves in newly-duplicated two-component systems (Capra *et al*, 2012; Laub & Goulian, 2007). NtrC-like activators and other bacterial DNA-binding response regulators have high affinity interactions with their cognate kinases, and crosstalk generally increases noise and decreases the overall response of the system to the incoming signal (Rowland & Deeds, 2014). The specificity of response regulators for phosphorylation by their cognate kinases is governed primarily by molecular recognition, though these proteins can be very sequence similar, and by maintaining a relatively high abundance of response regulator relative to its cognate kinase within a cell to prevent unwanted phosphorylation (Skerker *et al*, 2008; Capra *et al*, 2012; Podgornaia & Laub, 2013). Taken together, this indicates that mutations to response regulators, like those that we have introduced in the NtrC-like proteins in this study, might lead to a situation where there is a high concentration of phosphorylated kinase in the absence of its highest affinity interaction partner, allowing the cognate kinase to phosphorylate structurally similar non-target response regulators and inappropriately initiate those signaling cascades. This model would explain why so many of the NtrC-like activator mutations produced severe phenotypes that fail to form fruiting bodies in the same way and highlights that redundancy due to gene duplication can have negative consequences without proper insulation.

Rarely is a typical phenotype expressed a majority of the time within knockouts of homologous group – instead, the typical phenotype represents the plurality behavior. Replicates even of the same strain are seen to straddle multiple zones in the PCA feature space. This is particularly evident in the One component gene family, which was observed to have the greatest variation in phenotype from replicate to replicate (Fig. S2). The same mutant strain, for example, can yield a phenotype indistinguishable from wild-type in one replicate, and make immature aggregates or fail to aggregate entirely in other replicates. This variability might indicate that the impact of mutation on these functional networks increases sensitivity to stochastic fluctuations within the cellular environment that contribute to a tendency toward one phenotypic fate or another.

We do not propose that phenotypic similarity serves as a strong indicator of functional redundancy. There are almost certainly insignificant associations in the PCA feature space. For example, there are a small number of ABC transporter mutants positioned within the NtrC-like activator cluster. We do not propose that these genes are functionally redundant with the majority of NtrC-like activators, but we might suggest that they are less likely to be functionally redundant with those in the ABC transporter cluster. It is also possible that any group of completely unrelated genes could have some degree of functional redundancy, but this represents a background level or lower threshold of observable redundancy. We have shown that the redundancy we observe is significantly above that background by measuring the phenotypic clustering of random groupings of genes from the various homologous groups. We are sure there are many forms and many degrees of functional redundancy that are not represented by this PCA, but it does reveal a widely distributed functional redundancy above a background threshold.

Extensive progress has been made in recent years in looking at large-scale studies of digenic and even trigenic interactions and redundancies that affect phenotypic robustness and fitness (Kuzmin *et al*, 2021; Gagarinova *et al*, 2012). Others have begun to disentangle the relationship between subsets of multigene families and their roles in redundancy (Johnstun *et al*, 2021). We add to this growing body of literature by exploring functional redundancy in large, homologous gene families of *M. xanthus*. We make no claims about functional redundancy between specific genes in *M. xanthus*. Rather, we seek to define the scale and distributive nature of redundancy networks that include these large gene families, demonstrating that redundancy networks are not necessarily limited to a few of the closest paralogs; they may include dozens or hundreds of genes.

Instead of trying to quantify the direct effect of mutations on fitness by measuring a single variable such as growth, we chose to measure multiple aspects of a complex development process. While our method requires the collection of more data per strain than a synthetic genetic array, for example, it has the ability to detect more subtle phenotypes that may not have strong implications for fitness but can still inform studies of redundancy. Since many single gene disruptions have such subtle phenotypes, and since we propose that extensive redundancy networks protect an organism from the fitness costs of mutation, we chose phenotypic similarity, rather than overall effect on fitness, to assess the extent of functional redundancy within gene families.

Our results highlight the importance of considering the nature and extent of redundancy when making claims regarding interactions between genotype and phenotype. Gene families can have high degrees of functional entanglement that may mitigate the impact of mutation, so that quantifying even minor deviations in phenotype may allow for the recognition of patterns; if mutations within a redundancy network produce similar phenotypes, then subtle changes in phenotype have the potential to inform annotation. For example, a gene of unknown function displaying a subtle phenotype similar to that of genes of known function could provide evidence that the unknown gene is part of a redundancy network. Our image analysis pipeline can be extended to future studies of *M. xanthus*, even under differing experimental conditions, for automated extraction of phenotypic features. Further, our dataset can be used to probe whether there are patterns in amino acid sequence homology that lead to functional redundancy by comparing the sequences of genes that are located within the family cluster on the PCA to those that are located outside the cluster and are presumably non-redundant. Underscoring all these results is the observation that without a sufficiently large collection of mutants and replicates, functional redundancy does not present itself clearly enough to be recognized.

## Conclusions

This work provides evidence for the existence of large networks of redundant genes as a means by which an organism such as *Myxococcus xanthus* can execute complex multicellular social behaviors robust to perturbations to gene function. We observe subtle deviations in phenotype, a distinct set for each homologous gene family, that present when knocking out any one gene within these redundancy groups. These subtle deviations are measurable due to the large number of time series included in our full dataset and the quantitative detail of the extracted phenotypic information, which in combination necessitate the automated analysis pipeline we have developed.

## Methods

### Strains and Culture Conditions

*Myxococcus xanthus* strain DK1622 was used as the wild-type for this study. All 265 mutant strains in the ABC Transporter, ECF sigma factor, NtrC-like activator, and One Component Signal Transduction System families (Table S3) were created using plasmid insertion via homologous recombination as previously described (Caberoy *et al*, 2003; Plamann *et al*, 1995) and modified by Yan et al, 2014. Briefly, 400-600bp internal fragments of each gene were PCR amplified and ligated into pCR^®^2.1-TOPO [Invitrogen]. The plasmids were amplified in *E. coli* before isolation and electroporation into *M. xanthus* DK1622, where the plasmid incorporates into the *M. xanthus* genome via the homologous region on the plasmid. PCR verification was used to confirm the location of each insertion.

Cells were grown overnight in CTTYE (1% Casein Peptone (Remel, San Diego, CA, USA), 0.5% Bacto Yeast Extract (BD Biosciences, Franklin Lakes, NJ, USA), 10 mM Tris (pH 8.0), 1 mM KH(H_2_)PO_4_, 8 mM MgSO_4_) with vigorous shaking at 32°C. Cultures of mutant strains were supplemented with 40µg/mL kanamycin. Cells were centrifuged to remove the nutrient broth, washed in TPM buffer (10 mM Tris (pH 7.6), 1 mM KH(H_2_)PO_4_, 8 mM MgSO_4_), and resuspended to a final concentration of 5×10^9^ cells/mL. For development assays, approximately 2.5×10^7^ cells were spotted onto TPM agar slide complexes, as previously described (Taylor & Welch, 2010).

### Imaging

Development assays for wild-type and mutant strains were carried out on TPM starvation agar slide complexes for 24 hours, with approximately three replicates per strain. Though it can take multiple days for cells within fruiting bodies to fully differentiate into spores, we generated time series of only the first 24 hours of development because wild-type cells show little to no observable change in fruiting body morphology, count, or behavior following this period at the magnification used. Time-lapse grayscale images were captured every 60 seconds under 4x magnification with a Nikon Eclipse E-400 microscope [Nikon Instruments] and SPOT Insight camera. ImageJ was used for processing the .TIFF images into time series for analysis.

### Multidimensional scaling of gene sequence dissimilarity

Amino acid sequences for the four homologous families were retrieved from NCBI and imported into the Multiple Sequence Alignment tool in Clustal Omega (Madeira *et al*, 2022), generating a percent identity matrix for all 265 proteins. This was then converted to a percent dissimilarity matrix and used as the input for the Classical Multidimensional Scaling package in R to generate plotting coordinates in two dimensions. Then Gaussian kernel density estimation was used to plot an estimate of the probability distribution function (plotted with increased opacity to represent higher probability) to guide the eye in identifying sub-clusters of similar genes within each paralogous group.

### Manual phenotyping

Manual preliminary phenotyping of the mutant strains in this study was performed using the time series described above. We will refer to mature aggregates as fruiting bodies for simplicity, though we did not test sporulation efficiency in this study. First, strains that failed to produce fruiting bodies at all within 24 hours across all replicates were labeled “no aggregation” mutants. Strains that formed initial aggregates that disassociated completely before the 24-hour mark were labeled as “fall apart”. Some strains, labeled “aggregate-reaggregate”, formed aggregates that initially fell apart, but new aggregates were formed that persisted and looked similar to wild-type by the endpoint of the time series.

To qualitatively determine the start time of aggregation, the time series were observed in sliding windows of 25 minutes to identify the window where initial aggregates were first formed. The average start time of 22 wild-type replicates was used for comparison, and mutant strains that had start times outside of one standard deviation of the mean of wild-type were considered either “early” or “late” aggregation mutants.

Wild-type fruiting bodies at 24 hours appear almost black in color and are roughly circular in brightfield images. Any strains that appeared to have these characteristics and initiated aggregation within the same window as wild-type were classified like-wild-type (LWT). Strains that initiated aggregation at a normal time but didn’t develop aggregates that were as dark in gray value as wild-type were labeled as “immature aggregation” mutants. Finally, mutants that did not display consistent phenotypes across replicates were classified as “variable”. A table of all mutant strains used in the study, as well as their manually assigned phenotype, can be found in Table S3.

### Automated phenotyping

Phenotype was automatically quantified for the mutant strains in this study by running 144 individual .TIFF images (ten minutes between each frame over 24 hours of total development) from each time series through a custom Python image processing and analysis pipeline to identify in each frame which pixels could belong to a fruiting body, based on their gray value. The information for the position and geometry of each aggregate was filtered to remove noise and spurious aggregates. This information was then collected for the entire time series to track individual fruiting bodies over time, revealing their fate and dynamics. This detailed data summary for each time series then had a list of eighteen specific numbers extracted from it, each of which captures one overall feature, such as average growth rate or the average size of final fruiting bodies. The values of these eighteen metrics together (a phenotypic vector) constitute the phenotype profile for that time series. The full details of the image processing pipeline and all phenotypic metrics are available in the SI.

A selection of 133 mutant time series were chosen at random from each paralogous group so as not to weigh any paralogous group more than the other. The phenotypic vector for each time series was calculated, and the values of each metric were shifted by a constant amount and scaled by a constant factor so that across the dataset, each metric had a mean of zero and a variance of one. This ensured that one metric would not supersede the others simply due to the magnitude of its units. PCA was performed on this normalized dataset to extract the two combinations of metrics, PC1 and PC2, that captured the most variation across the dataset.

The phenotypic clusters were revealed by plotting each time series as a point in the PC1 vs. PC2 phenotype space and then estimating the probability density for each homologous group via Gaussian kernel density estimation. Essentially, a Gaussian blur was applied to the points, and areas of greater overlap were colored with higher opacity, as shown in Fig. 4. The width of the smoothing kernel was chosen to be the smallest value that could preserve the shape of the probability density for different equally sized subsamples from each homologous group.

The statistical test used to generate the p-values for average cluster separation and average cluster size was a form of bootstrapping which started with the PC1 and PC2 coordinates of each point shown in the data sample of Fig. 4. Each point was randomly reassigned one of four arbitrary families in such a way that replicates of the same strain were all assigned the same family. A new Gaussian kernel density estimation was performed to approximate the probability density of each family in PCA phenotype space. The contour representing 75% of the maximum value of the estimated probability distribution function was then extracted, with cluster separation being the average across all pairings of families of the centroid-to-centroid distance between contours, and cluster size quantified by the average across families of the radius of gyration of each contour, i.e. the root mean square distance of each contour’s points from its centroid. Each p-value was calculated as the fraction of the random groupings that had a greater average separation or smaller average size than that of the original data grouped by the actual gene families.

## Acknowledgements

The authors acknowledge Linnea Ritchie and Christina McKeown for helpful comments and edits.

## Author Contributions

Conceptualization, experimental design, and data collection: MEA, FAB, JAC, AEP, and RDW; data analysis and validation: MEA, JAC, IL; writing and revision: MEA, JAC, AEP, RDW.

## References

Abellón-Ruiz J, Bernal-Bernal D, Abellán M, Fontes M, Padmanabhan S, Murillo FJ & Elías-Arnanz M (2014) The CarD/CarG regulatory complex is required for the action of several members of the large set of Myxococcus xanthus extracytoplasmic function σ factors. Environ Microbiol 16: 2475–2490

Baker EA, Gilbert SPR, Shimeld SM & Woollard A (2021) Extensive non-redundancy in a recently duplicated developmental gene family. BMC Ecol Evol 21: 1–17

Bretl DJ & Kirby JR (2016) Molecular Mechanisms of Signaling in Myxococcus xanthus Development. J Mol Biol 428: 3805–3830

Butland G, Babu M, Díaz-Mejía JJ, Bohdana F, Phanse S, Gold B, Yang W, Li J, Gagarinova AG, Pogoutse O, et al (2008) eSGA: E. coli synthetic genetic array analysis. Nat Methods 5: 789–795

Caberoy NB, Welch RD, Jakobsen JS, Slater SC & Garza AG (2003) Global Mutational Analysis of NtrC-Like Activators in Myxococcus xanthus: Identifying Activator Mutants Defective for Motility and Fruiting Body Development. J Bacteriol 185: 6083–6094

Capra EJ, Perchuk BS, Skerker JM & Laub MT (2012) Adaptive mutations that prevent crosstalk enable the expansion of paralogous signaling protein families. Cell 150: 222–232

Dean EJ, Davis JC, Davis RW & Petrov DA (2008) Pervasive and persistent redundancy among duplicated genes in yeast. PLoS Genet 4

diCenzo GC & Finan TM (2015) Genetic redundancy is prevalent within the 6.7 Mb Sinorhizobium meliloti genome. Mol Genet Genomics 290: 1345–1356

Diss G, Ascencio D, Deluna A & Landry CR (2014) Molecular mechanisms of paralogous compensation and the robustness of cellular networks. J Exp Zool Part B Mol Dev Evol 322: 488–499

Durmort C & Brown JS (2015) Streptococcus pneumoniae Lipoproteins and ABC Transporters. Streptococcus Pneumoniae Mol Mech Host-Pathogen Interact: 181–206

Gagarinova A, Babu M, Greenblatt J & Emili A (2012) Mapping bacterial functional networks and pathways in Escherichia coli using synthetic genetic arrays. J Vis Exp: 1–11

Giaever G, Chu AM, Ni L, Connelly C, Riles L, Veronneau S, Dow S, Lucau-danila A, Anderson K, Arkin AP, et al (2002) Functional profiling of the Saccharomyces cerevisiae genome. Nature: 387–391

Goldman BS, Nierman WC, Kaiser D, Slater SC, Durkin AS, Eisen JA, Monning CM, Barbazuk WB, Blanchard M, Field C, et al (2006) Evolution of sensory complexity recorded in a myxobacterial genome. Proc Natl Acad Sci U S A 103: 15200–15205

Hannay K, Marcotte EM & Vogel C (2008) Buffering by gene duplicates: An analysis of molecular correlates and evolutionary conservation. BMC Genomics 9

Johnstun JA, Shankar V, Mokashi SS, Sunkara LT, Ihearahu UE, Lyman RL, Mackay TFC & Anholt RRH (2021) Functional Diversification, Redundancy, and Epistasis among Paralogs of the Drosophila melanogaster Obp50a-d Gene Cluster. Mol Biol Evol 38: 2030–2044

Krakauer DC & Plotkin JB (2002) Redundancy, antiredundancy, and the robustness of genomes. Proc Natl Acad Sci U S A 99: 1405–1409

Kuzmin E, Rahman M, VanderSluis B, Costanzo M, Myers CL, Andrews BJ & Boone C (2021) τ-SGA: synthetic genetic array analysis for systematically screening and quantifying trigenic interactions in yeast. Nat Protoc 16: 1219–1250

Kuzmin E, Taylor JS & Boone C (2022) Retention of duplicated genes in evolution. Trends Genet 38: 59–72

Laub MT & Goulian M (2007) Specificity in two-component signal transduction pathways. Annu Rev Genet 41: 121–145

Lehár J, Krueger A, Zimmermann G & Borisy A (2008) High-order combination effects and biological robustness. Mol Syst Biol 4: 1–6

Luo Y & Helmann JD (2009) Extracytoplasmic function σ factors with overlapping promoter specificity regulate sublancin production in Bacillus subtilis. J Bacteriol 191: 4951–4958

Madeira F, Pearce M, Tivey ARN, Basutkar P, Lee J, Edbali O, Madhusoodanan N, Kolesnikov A & Lopez R (2022) Search and sequence analysis tools services from EMBL-EBI in 2022. Nucleic Acids Res: 1–4

Mascher T, Hachmann AB & Helmann JD (2007) Regulatory overlap and functional redundancy among Bacillus subtilis extracytoplasmic function σ factors. J Bacteriol 189: 6919–6927

Ohno S (1970) Evolution by Gene Duplication Springer Berlin Heidelberg

Orelle C, Mathieu K & Jault JM (2019) Multidrug ABC transporters in bacteria. Res Microbiol 170: 381–391

Paget MS (2015) Bacterial sigma factors and anti-sigma factors: Structure, function and distribution. Biomolecules 5: 1245–1265

Plamann L, Li Y, Cantwell B & Mayor J (1995) The Myxococcus xanthus asgA gene encodes a novel signal transduction protein required for multicellular development. J Bacteriol 177: 2014–2020

Podgornaia AI & Laub MT (2013) Determinants of specificity in two-component signal transduction. Curr Opin Microbiol 16: 156–162

Rees DC, Johnson E & Lewinson O (2009) ABC transporters: the power to change. Nat Rev Mol Cell Biol 10: 218–27

Rhodius VA, Segall-Shapiro TH, Sharon BD, Ghodasara A, Orlova E, Tabakh H, Burkhardt DH, Clancy K, Peterson TC, Gross CA, et al (2013) Design of orthogonal genetic switches based on a crosstalk map of σs, anti-σs, and promoters. Mol Syst Biol 9: 1–13

Rowland MA & Deeds EJ (2014) Crosstalk and the evolution of specificity in two-component signaling. Proc Natl Acad Sci U S A 111: 5550–5555

Skerker JM, Perchuk BS, Siryaporn A, Lubin EA, Ashenberg O, Goulian M & Laub MT (2008) Rewiring the Specificity of Two-Component Signal Transduction Systems. Cell 133: 1043–1054

Taylor RG & Welch RD (2010) Recording Multicellular Behavior in Myxococcus xanthus Biofilms using Time-lapse Microcinematography. J Vis Exp: 1–6

Thomaides HB, Davison EJ, Burston L, Johnson H, Brown DR, Hunt AC, Errington J & Czaplewski L (2007) Essential bacterial functions encoded by gene pairs. J Bacteriol 189: 591–602

Vandersluis B, Bellay J, Musso G, Costanzo M, Papp B, Vizeacoumar FJ, Baryshnikova A, Andrews B, Boone C & Myers CL (2010) Genetic interactions reveal the evolutionary trajectories of duplicate genes. Mol Syst Biol 6: 1–13

Wagner A & Wright J (2007) Alternative routes and mutational robustness in complex regulatory networks. BioSystems 88: 163–172

Xie C, Zhang H, Shimkets LJ & Igoshin OA (2011) Statistical image analysis reveals features affecting fates of Myxococcus xanthus developmental aggregates. Proc Natl Acad Sci U S A 108: 5915–20

Yan J, Bradley MD, Friedman J & Welch RD (2014) Phenotypic profiling of ABC transporter coding genes in Myxococcus xanthus. Front Microbiol 5: 1–12

