## Supplementary Information for "Phenotypic similarity is a measure of functional redundancy within homologous gene families"

Jessica A. Comstock, Merrill E. Asp, Fatmagül Bahar, Isabella Lee, Alison E. Patteson, Roy D. Welch

*Statistical testing and controls*

The results presented in Fig. 4, namely the separation of phenotypic clusters for each homologous gene family, were tested for statistical significance by recreating the estimated probability density functions for random groupings instead of the actual gene family and observing the resulting sharpness of and separation between peaks in each grouping’s probability density function (see Methods). To illustrate the difference in phenotypic peak between these random groupings and the actual gene family groupings expected to correspond to redundant gene groups, we present the results of a representative random grouping in Fig. S1.

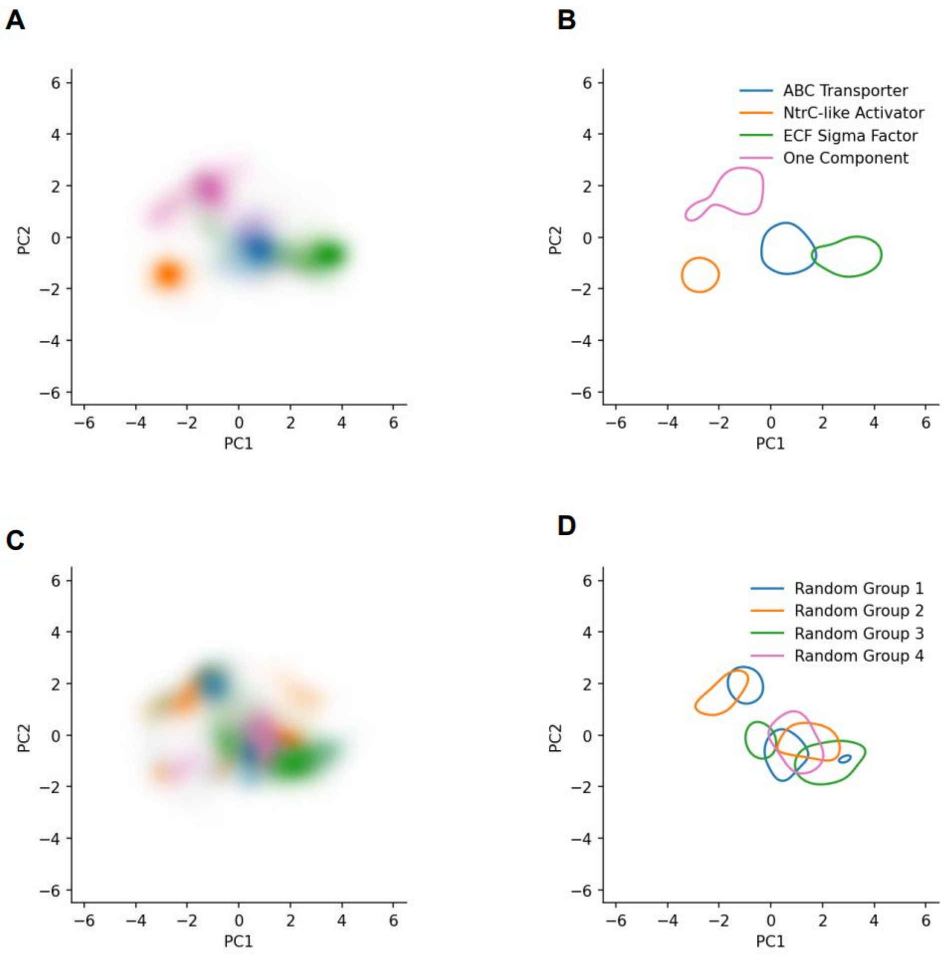


**Figure S1**: Phenotypic clusters arise robustly from homologous gene families as compared to random groupings of mutant strains. **(A)** Reproduced from Figure 4, each of the four gene families produces a distinct phenotypic cluster when plotting the estimated probability distribution function for that family (using Gaussian kernel density estimation) in phenotype space. **(B)** A contour is shown for each gene family where the estimated probability distribution function is at 75% of its maximum value, and the geometry of that contour is used to quantify the width of the cluster and its separation from other clusters. **(C)** The same data used in Figure 4 was regrouped into four random groupings, and the PCA and probability density function estimates were repeated, showing much more incoherent phenotype clusters. **(D)** The corresponding contours for the four random groupings show less separation and less sharpness than phenotypic clusters based on homologous gene groups. Both (C) and (D) come from one representative random grouping, many of which were made to calculate the p-values for cluster separation and sharpness reported in Results.

Replicates of the same strain can vary in phenotype. Several plots showing phenotypic spread for a few representative strains are included in Figure S2.

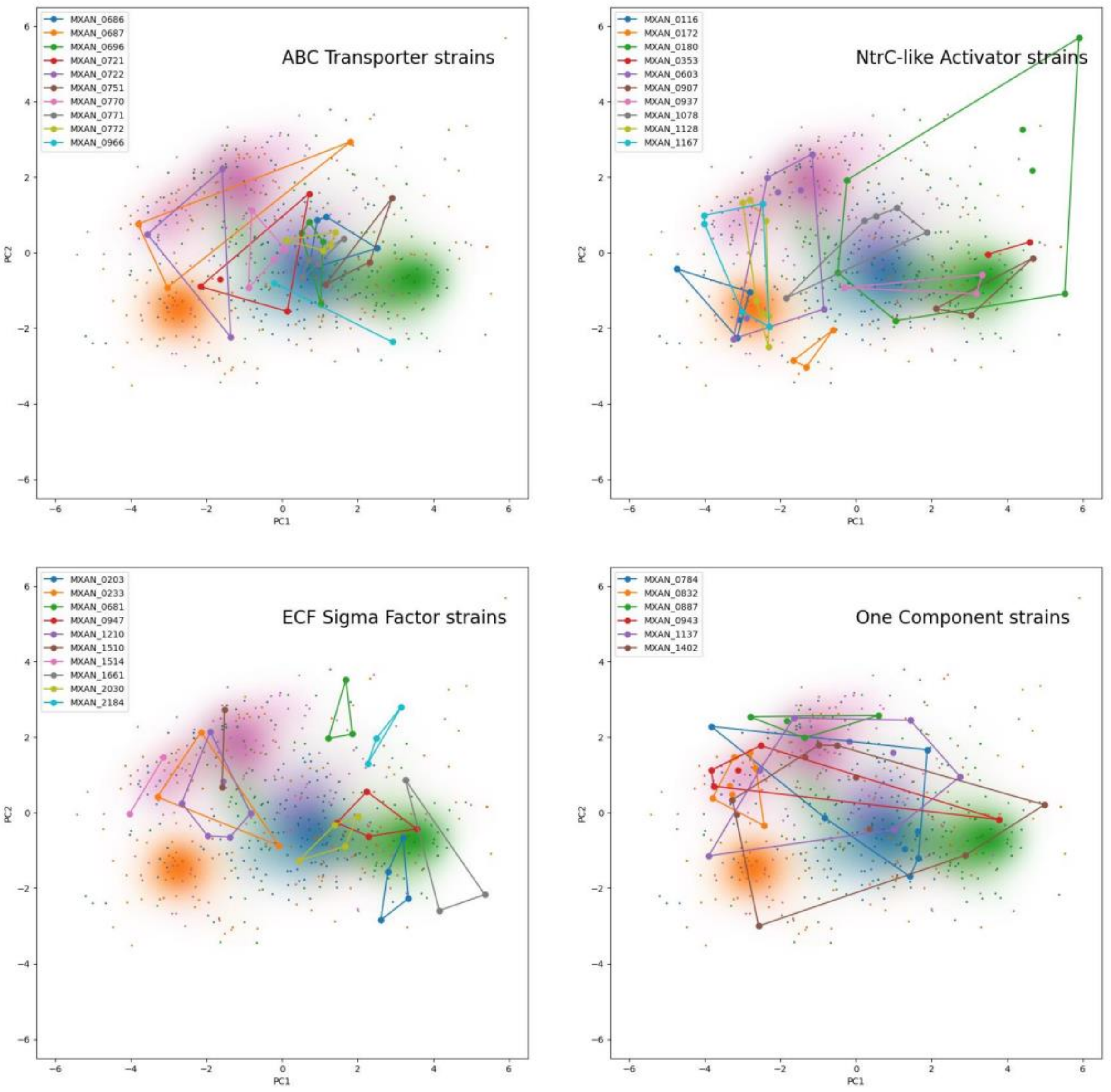


**Figure S2**: Replicates of the same strain can vary in phenotype. Reproduced for context from Figure 4 are the phenotypic scatterplot resulting from the PCA (where each point is a time series, plotted nearby other time series that are phenotypically similar) and the superimposed probability distribution functions for each of the homologous gene families: ABC transporters in blue, NtrC-like activators in orange, ECF sigma factors in green, and One component in pink. Each subplot includes all replicates of a few representative strains, where the replicates of each strain are represented in a single color and drawn with a bounding polygon to aid the eye. Replicate-to-replicate variation is larger or smaller depending on strain and to which homologous family the strain belongs.

A metric for replicate-to-replicate phenotypic spread of a specific strain is the sample standard deviation $s$ generalized to two dimensions

$$s = \sqrt{\frac{1}{n-1}\sum_{i=1}^{n}((x_i - \bar{x})^2 + (y_i - \bar{y})^2)}$$

where $n$ is the number of replicate points, $x_i$ and $y_i$ are the coordinates of the $i$ th replicate point, and $\bar{x}$ and $\bar{y}$ are the means of each coordinate across the replicate points, i.e. the centroid coordinates. In this case, the $x$ and $y$ coordinates are the value of PC1 and PC2 respectively.

Table S1 summarizes the mean replicate-to-replicate phenotype spread averaged over strain for each gene family using the metric $s$, with errors given by the standard error of the means.

| **Gene family** | **Mean replicate-to-replicate spread $s$** |
|---|---|
| ABC Transporters | 1.42 ± 0.12 |
| NtrC-like Activators | 1.57 ± 0.14 |
| ECF Sigma Factors | 1.31 ± 0.09 |
| One Component | 1.94 ± 0.18 |

**Table S1**: A summary of the average replicate-to-replicate spread for each homologous gene family, with errors given by the standard error of the means. This spread is illustrated for some representative strains in Fig. S2

This indicates a statistically significant difference for replicate-to-replicate phenotype spread between One Component strains and ABC Transporter strains (p = 0.024), and between One Component strains and ECF Sigma Factor strains (p = 0.004) according to a two-sided Welch's t-test.

*Correlation of genetic differences with phenotypic differences*

When comparing the genetic similarity and phenotypic similarity of the mutant strains used in this study, little correlation was found, as illustrated in Figure S3.

For each homologous gene group, all unique pairings of different strains were plotted using genetic difference on the x-axis and phenotypic difference on the y-axis. Genetic difference was quantified using Clustal Omega Multiple Sequence Alignment (Madeira *et al*, 2022), and phenotypic difference was calculated as the Euclidean distance between two 18-feature points, averaging the feature values over all of the replicates for that strain. These plots show that close genetic similarity of two mutant strains is not necessary for and in fact poorly predicts phenotypic similarity. Instead, we infer that phenotypic similarity is roughly equivalent across all mutants in a group of redundant genes.

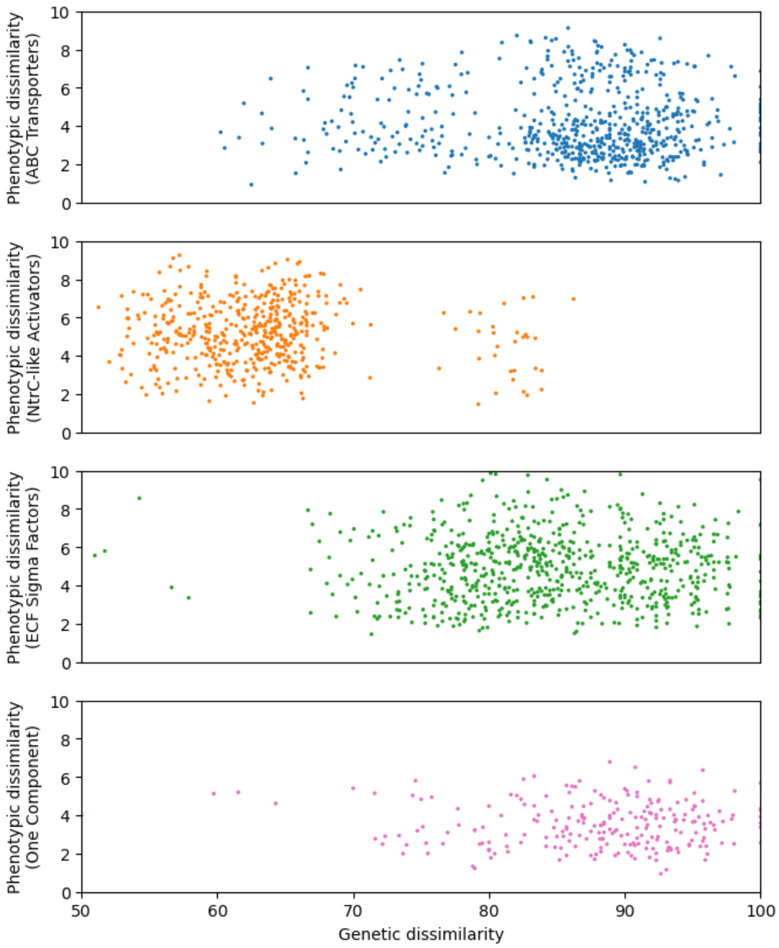


**Figure S3**: Genetic similarity is not an effective predictor of phenotypic similarity within a homologous gene family. For each of the four gene families analyzed, each point represents a unique pairing of two strains. Phenotypic dissimilarity is quantified by Euclidean distance in 18-dimensional feature space, where feature values are represented by averages over all replicates for that strain. Genetic dissimilarity is quantified by comparison of base pairs using Clustal Omega Multiple Sequence Alignment. Within each of the four gene families, phenotypic dissimilarity and genetic dissimilarity to not correlate.

*Description of typical phenotype for each homologous gene family*

A manual review of time series near the phenotypic cluster for each respective gene family (Fig. 4) was performed to describe the typical behavior. This summary is with respect to only time series that fell within the contour of 75% of the maximum of the probability distribution function for each gene family.

Both the ABC transporter and ECF sigma factor mutants showed similar behaviors both with rapid formation and darkening of aggregates. Their differences lie in the number of persistent fruiting bodies and the rates of formation. Fruiting bodies of ECF sigma factor mutants form faster than the those of ABC transporter mutants, captured by growth rate.

The time series in the ECF sigma factor cluster also have a higher fraction of their fruiting bodies fail to persist. So, although they are fast to form, these aggregates have more of a tendency to either be absorbed by their neighboring fruiting bodies or not survive at all.

Nearly all of the time series in the NtrC-like activator cluster fail to form fruiting bodies. While the bacteria do move around and occasionally form aggregates, these never completely darken or stabilize and most evaporate. This tendency to fail to aggregate is most directly captured quantitatively in the small final number of persistent fruiting bodies and in the low growth rate of these time series.

One component mutants near the phenotypic cluster were observed to have the ability to form fruiting bodies but not darken significantly, remaining immature. They also appear to be less circular than the fruiting bodies of ABC transporters or ECF sigma factor mutants. The One component time series also showed signs of struggle when forming – they took much longer than the other successfully aggregating gene families to form their persistent fruiting bodies.

The individual phenotypic features that support these observations are included in Fig. S4.

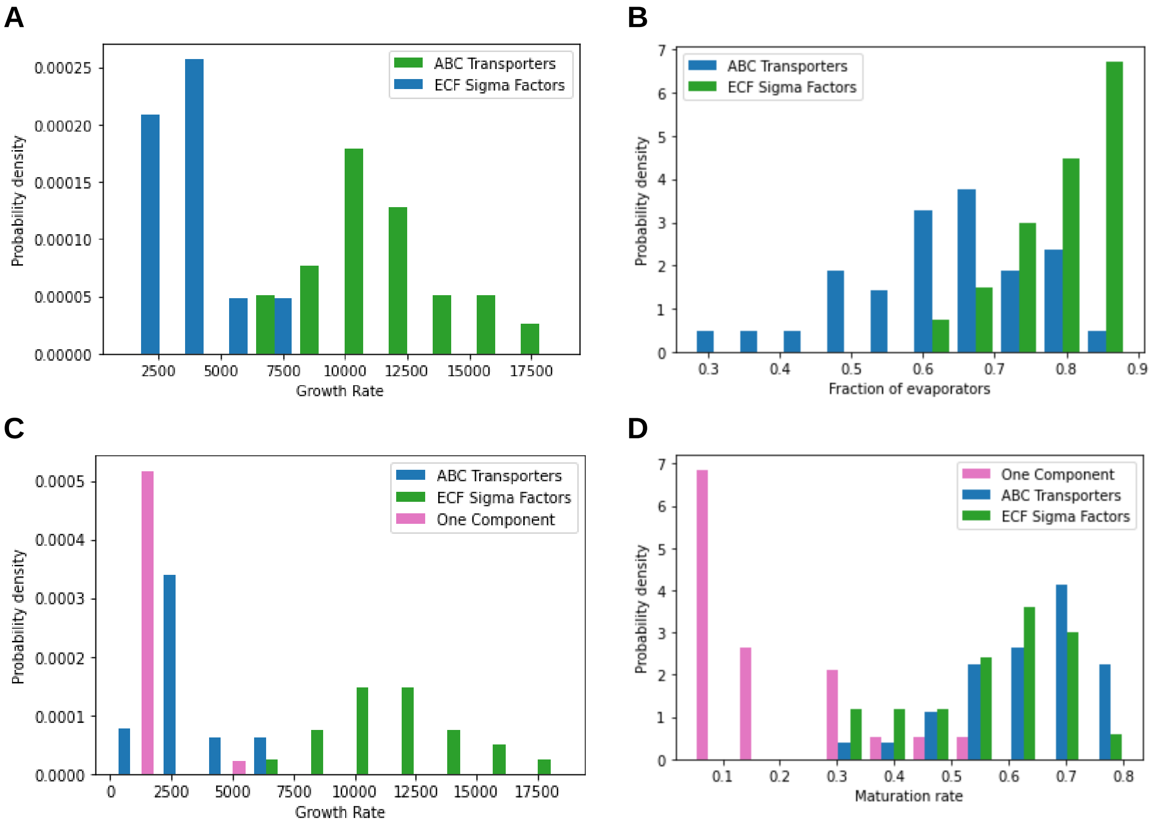


**Figure S4**: Quantitative comparison of the behavior in phenotypic clusters for each gene family. **(A)** Histogram of growth rates. ECF sigma factor mutants form aggregates faster than ABC transporter mutants. **(B)** Histogram of fraction of fruiting bodies that fail to persist. ECF sigma factor mutants have more evaporating fruiting bodies than ABC transporter mutants. **(C)** Histogram of growth rates between three gene families with successful fruiting body formation. One component mutants form fruiting bodies the slowest of the three displayed gene families. **(D)** Histogram of maturation rates (i.e. the maximum rate of darkening during fruiting body maturation). One component mutants do not darken or mature at a rate much slower than ABC transporter mutants or ECF sigma factor mutants.

*Image processing*

Custom Python code was written for this analysis, available on Github (https://github.com/masp01/SU-myxo-aggregate-tracking). Using the Python implementation of OpenCV (Bradski, 2000), each raw frame is put through the following image processing steps to identify the size, shape, and position of each fruiting body:

1. Non-local means denoising (cv2.fastNlMeansDenoising)
   To remove background noise, a 7 pixel wide template window is moved over the image to find regions that visually match (typically uniform, noisy regions). This search is done within a 21 pixel distance of each patch of the image. The gray value of each pixel is replaced with the average gray value of pixels in matching regions, smoothing over noise while keeping boundaries distinct. The smoothing strength was chosen at a constant value of 70 after manually testing parameter values for many images. Template window and search sizes are standard and were not tuned.
2. Adaptive thresholding (cv2.adaptiveThreshold)
   To identify locally dark regions that should belong to fruiting bodies, the gray value of each pixel is compared to the average gray value of its neighbors within a block 101 pixels (145 µm) wide. 8-bit pixels (gray value from 0 – 255) that have a gray value at least 20 below this local average are marked white. All other pixels are marked black, creating a binary image. Parameters were chosen after manually testing with many images and are robust enough to be used across the entire dataset.
3. Morphological opening (cv2.morphologyEx)
   A circular kernel 5 pixels (7.2 µm) wide is moved over the binary image. Any feature covered entirely by the kernel is removed. This reduces single-pixel noise.
4. Contour identification (cv2.findContours)
   Contiguous regions of white pixels are automatically identified in the binary image. A list is compiled of the x,y coordinates of the pixels on the boundary of each such region. This gives both a count of total candidate fruiting bodies and the geometry of their boundary.

At this point, a list of features has been identified, some of which are genuine fruiting bodies, and some of which are noise or spurious aggregates. The contour of each feature is measured for the x,y coordinates of its center, its area $A$, perimeter $P$, and average gray value. The circularity $4\pi A/P^2$, is also calculated. It captures the elongation of the fruiting bodies and ranges from 0 (completely flat) to 1 (perfectly circular).

*Tracking fruiting bodies and filtering*

Once all the frames of a time series have been processed, the Python package Trackpy (Allan, D. B., Caswell, T., Keim, N. C. & van der Wel, C. M. *trackpy: Trackpy v0.4.2* doi:10.5281/zenodo.4682814). is used to assign an ID to each feature that tracks it over time. It is at this point that filtering is done to remove spurious features:

1. Minimum area filter
   Features that are smaller than 576 µm² are ignored. This is the smallest fruiting body size that is distinguishable from noise at 4X magnification

2. Maximum gray value filter
   Features with an average gray value above 200 (max 255) are considered too bright to be fruiting bodies and are ignored.
3. Formation time filter
   Features that appear before 100 minutes have elapsed are incidental initial aggregates, and not genuine fruiting bodies that have assembled over time. In no time series did a new aggregate form in less than 100 minutes. These incidental aggregates are tracked over time and ignored in all frames in which they appear.
4. Category filter
   The area dynamics of each remaining fruiting body are considered to see if the fruiting body persists to the end of the time series (persistors) or if it vanishes smoothly (evaporators). Persistors with an average circularity below 0.5 are typically noise and are ignored. Smoothly vanishing is defined as starting with an area less than max area and then decreasing from maximum area by at least 25% by the final frame of the time series. Evaporators with centers within (14.4 µm of the edge of the frame or with an average circularity below 0.5 are considered noise and ignored. Any feature that cannot be categorized as a persistor or evaporator is assumed to be spurious or contain dynamics errors and is ignored.

*Feature extraction*

The data for each time series is then analyzed to measure the following quantitative features, each a single number summarizing one aspect of the time series. Measurements are taken over a 7.2 $mm^2$ field size.

**Table S2**: Enumeration of all quantitative features used in the automated phenotype analysis

| | Feature name | Description | Formula |
|---|---|---|---|
| 1 | Start time | Elapsed time from inoculation to the beginning of visible aggregation | When at least 10 fruiting bodies grow larger than 1000 $\mu m^2$ in area |
| 2 | Peak time | Elapsed time from inoculation to the peak of visible aggregation | When total fruiting body area reaches a maximum |
| 3 | Stability time | Elapsed time from inoculation to overall aggregate stability | When the number of fruiting bodies changes by less than three per hour (24 hours maximum) |
| 4 | Growth time | Duration of initial growth phase | Peak time minus start time |
| 5 | Growth rate | Average rate of total area increase during growth phase | Change in total area divided by change in time between start time and peak time |
| 6 | Peak average area | The average fruiting body area at peak time | |

| | | | |
|---|---|---|---|
| 7 | Peak area std | The standard deviation of fruiting body area at peak time | |
| 8 | Final average area | The average fruiting body area at the moment 24 hours after inoculation | |
| 9 | Final area std | The standard deviation of fruiting body area at the moment 24 hours after inoculation | |
| 10 | Gray value % change | Percent difference between minimum and maximum average gray value (only for persistent fruiting bodies) | |
| 11 | Maturation rate | Maximum slope of gray value vs. time curve for persistent fruiting bodies | |
| 12 | Temporal coherence | How closely in time each evaporating fruiting body reaches its maximum area before starting to shrink | The standard deviation of the distribution of the time of maximum area for evaporators |
| 13 | Fraction of evaporators | Total number of evaporators divided by total number of evaporators plus persistors | |
| 14 | Maximum number | Total number of fruiting bodies at peak time | |
| 15 | Average lifetime | The elapsed time between an evaporator's first and final moment above the minimum area threshold, averaged over all evaporators | |
| 16 | Std lifetime | Standard deviation of the elapsed times between each evaporator's first and final moment above the minimum area threshold | |
| 17 | Maximum average area falloff | | Most negative slope of average area vs. time curve |
| 18 | Maximum number falloff | | Most negative slope of number vs. time curve |

**Table S3**: List of strains used in this study, listed by MXAN number, followed by the manual phenotype classification associated with Figure 2, gene family, and citation for creation of the specific strain used.

| **Strain** | **Phenotypic Classification** | **Gene Family** | **Strain Creation** |
|---|---|---|---|
| DK1622 | Wild-type | N/A | |
| MXAN_0035 | No aggregation | ABC Transporter | (Yan *et al*, 2014) |
| MXAN_0036 | LWT | ABC Transporter | (Yan *et al,* 2014) |
| MXAN_0037 | Late aggregation | ABC Transporter | (Yan *et al,* 2014) |
| MXAN_0069 | Early aggregation | One Component | This study |
| MXAN_0079 | Variable | One Component | (Ritchie *et al*, 2021) |
| MXAN_0090 | Late aggregation | One Component | This study |
| MXAN_0107 | Early aggregation | ABC Transporter | (Yan *et al,* 2014) |
| MXAN_0108 | LWT | ABC Transporter | (Yan *et al,* 2014) |
| MXAN_0116 | No aggregation | NtrC-Like Activators | This study |
| MXAN_0172 | Fall apart | NtrC-Like Activators | (Ritchie *et al,* 2021) |
| MXAN_0180 | LWT | NtrC-Like Activators | This study |
| MXAN_0203 | Early aggregation | ECF Sigma Factors | This study |
| MXAN_0213 | Variable | One Component | (Ritchie *et al,* 2021) |
| MXAN_0214 | Variable | One Component | This study |
| MXAN_0233 | Immature aggregates | ECF Sigma Factors | This study |
| MXAN_0250 | Early aggregation | ABC Transporter | (Yan *et al,* 2014) |
| MXAN_0251 | LWT | ABC Transporter | (Yan *et al,* 2014) |
| MXAN_0353 | Early aggregation | NtrC-Like Activators | This study |
| MXAN_0387 | Late aggregation | One Component | This study |
| MXAN_0445 | Early aggregation | One Component | This study |
| MXAN_0502 | Variable | One Component | (Ritchie *et al,* 2021) |
| MXAN_0553 | LWT | ABC Transporter | (Yan *et al,* 2014) |
| MXAN_0554 | LWT | ABC Transporter | (Yan *et al,* 2014) |
| MXAN_0556 | LWT | One Component | This study |
| MXAN_0559 | Variable | ABC Transporter | (Yan *et al,* 2014) |
| MXAN_0596 | Immature aggregates | ABC Transporter | (Yan *et al,* 2014) |
| MXAN_0597 | LWT | ABC Transporter | (Yan *et al,* 2014) |
| MXAN_0603 | Late aggregation | NtrC-Like Activators | (Ritchie *et al,* 2021) |
| MXAN_0622 | LWT | ABC Transporter | (Yan *et al,* 2014) |
| MXAN_0627 | Early aggregation | One Component | This study |
| MXAN_0629 | Immature aggregates | ABC Transporter | (Yan *et al,* 2014) |
| MXAN_0654 | LWT | One Component | This study |
| MXAN_0665 | Variable | One Component | (Ritchie *et al,* 2021) |
| MXAN_0681 | Aggregate-reaggregate | ECF Sigma Factors | (Ritchie *et al,* 2021) |
| MXAN_0684 | Late aggregation | ABC Transporter | (Yan *et al,* 2014) |
| MXAN_0685 | No aggregation | ABC Transporter | (Yan *et al,* 2014) |

| | | | |
|---|---|---|---|
| MXAN_0686 | LWT | ABC Transporter | (Yan *et al*, 2014) |
| MXAN_0687 | Late aggregation | ABC Transporter | (Yan *et al*, 2014) |
| MXAN_0696 | Early aggregation | ABC Transporter | (Yan *et al*, 2014) |
| MXAN_0707 | Variable | One Component | This study |
| MXAN_0721 | LWT | ABC Transporter | (Yan *et al*, 2014) |
| MXAN_0722 | LWT | ABC Transporter | (Yan *et al*, 2014) |
| MXAN_0748 | LWT | One Component | This study |
| MXAN_0751 | LWT | ABC Transporter | (Yan *et al*, 2014) |
| MXAN_0770 | LWT | ABC Transporter | (Yan *et al*, 2014) |
| MXAN_0771 | LWT | ABC Transporter | (Yan *et al*, 2014) |
| MXAN_0772 | LWT | ABC Transporter | (Yan *et al*, 2014) |
| MXAN_0832 | Variable | One Component | This study |
| MXAN_0887 | Immature aggregates | One Component | This study |
| MXAN_0907 | Early aggregation | NtrC-Like Activators | This study |
| MXAN_0937 | LWT | NtrC-Like Activators | This study |
| MXAN_0943 | Variable | One Component | This study |
| MXAN_0947 | LWT | ECF Sigma Factors | This study |
| MXAN_0966 | Early aggregation | ABC Transporter | (Yan *et al*, 2014) |
| MXAN_0967 | LWT | ABC Transporter | (Yan *et al*, 2014) |
| MXAN_0968 | Late aggregation | ABC Transporter | (Yan *et al*, 2014) |
| MXAN_0995 | LWT | ABC Transporter | (Yan *et al*, 2014) |
| MXAN_1078 | Immature aggregates | NtrC-Like Activators | (Ritchie *et al*, 2021) |
| MXAN_1097 | LWT | ABC Transporter | (Yan *et al*, 2014) |
| MXAN_1124 | LWT | ABC Transporter | (Yan *et al*, 2014) |
| MXAN_1128 | Variable | NtrC-Like Activators | (Ritchie *et al*, 2021) |
| MXAN_1137 | Variable | One Component | This study |
| MXAN_1151 | Variable | ABC Transporter | (Yan *et al*, 2014) |
| MXAN_1153 | LWT | ABC Transporter | (Yan *et al*, 2014) |
| MXAN_1154 | LWT | ABC Transporter | (Yan *et al*, 2014) |
| MXAN_1155 | LWT | ABC Transporter | (Yan *et al*, 2014) |
| MXAN_1167 | Late aggregation | NtrC-Like Activators | (Ritchie *et al*, 2021) |
| MXAN_1189 | Early aggregation | NtrC-Like Activators | This study |
| MXAN_1210 | Late aggregation | ECF Sigma Factors | This study |
| MXAN_1245 | Immature aggregates | NtrC-Like Activators | (Ritchie *et al*, 2021) |
| MXAN_1262 | LWT | ABC Transporter | (Yan *et al*, 2014) |
| MXAN_1286 | Aggregate-reaggregate | ABC Transporter | (Yan *et al*, 2014) |
| MXAN_1319 | LWT | ABC Transporter | (Yan *et al*, 2014) |
| MXAN_1320 | Early aggregation | ABC Transporter | (Yan *et al*, 2014) |
| MXAN_1321 | Early aggregation | ABC Transporter | (Yan *et al*, 2014) |
| MXAN_1345 | LWT | NtrC-Like Activators | This study |
| MXAN_1376 | Late aggregation | ABC Transporter | (Yan *et al*, 2014) |

| | | | |
|---|---|---|---|
| MXAN_1377 | LWT | ABC Transporter | (Yan *et al*, 2014) |
| MXAN_1402 | Variable | One Component | This study |
| MXAN_1510 | LWT | ECF Sigma Factors | This study |
| MXAN_1514 | Immature aggregates | ECF Sigma Factors | This study |
| MXAN_1547 | LWT | ABC Transporter | (Yan *et al*, 2014) |
| MXAN_1548 | LWT | ABC Transporter | (Yan *et al*, 2014) |
| MXAN_1565 | Variable | NtrC-Like Activators | (Ritchie *et al*, 2021) |
| MXAN_1575 | Variable | One Component | This study |
| MXAN_1597 | LWT | ABC Transporter | (Yan *et al*, 2014) |
| MXAN_1598 | Variable | ABC Transporter | (Yan *et al*, 2014) |
| MXAN_1604 | Variable | ABC Transporter | (Yan *et al*, 2014) |
| MXAN_1605 | Early aggregation | ABC Transporter | (Yan *et al*, 2014) |
| MXAN_1654 | LWT | One Component | This study |
| MXAN_1661 | Early aggregation | ECF Sigma Factors | This study |
| MXAN_1667 | LWT | One Component | This study |
| MXAN_1677 | Variable | One Component | This study |
| MXAN_1683 | LWT | One Component | This study |
| MXAN_1695 | Immature aggregates | ABC Transporter | (Yan *et al*, 2014) |
| MXAN_1711 | No aggregation | One Component | This study |
| MXAN_1719 | LWT | One Component | This study |
| MXAN_1726 | Variable | One Component | This study |
| MXAN_1746 | Variable | One Component | This study |
| MXAN_1757 | Variable | One Component | This study |
| MXAN_2018 | LWT | ABC Transporter | (Yan *et al*, 2014) |
| MXAN_2019 | No aggregation | ABC Transporter | (Yan *et al*, 2014) |
| MXAN_2020 | Immature aggregates | ABC Transporter | (Yan *et al*, 2014) |
| MXAN_2030 | Early aggregation | ECF Sigma Factors | (Ritchie *et al*, 2021) |
| MXAN_2078 | LWT | ABC Transporter | (Yan *et al*, 2014) |
| MXAN_2128 | Immature aggregates | One Component | This study |
| MXAN_2145 | Late aggregation | One Component | This study |
| MXAN_2159 | Early aggregation | NtrC-Like Activators | This study |
| MXAN_2184 | Aggregate-reaggregate | ECF Sigma Factors | This study |
| MXAN_2204 | Immature aggregates | ECF Sigma Factors | This study |
| MXAN_2230 | Late aggregation | One Component | This study |
| MXAN_2234 | Immature aggregates | One Component | This study |
| MXAN_2249 | Late aggregation | ABC Transporter | (Yan *et al*, 2014) |
| MXAN_2250 | LWT | ABC Transporter | (Yan *et al*, 2014) |
| MXAN_2251 | Early aggregation | ABC Transporter | (Yan *et al*, 2014) |
| MXAN_2268 | LWT | ABC Transporter | (Yan *et al*, 2014) |
| MXAN_2395 | Early aggregation | ECF Sigma Factors | This study |
| MXAN_2407 | LWT | ABC Transporter | (Yan *et al*, 2014) |

| | | | |
|---|---|---|---|
| MXAN_2428 | LWT | ABC Transporter | (Yan *et al,* 2014) |
| MXAN_2429 | LWT | ABC Transporter | (Yan *et al,* 2014) |
| MXAN_2430 | LWT | ABC Transporter | (Yan *et al,* 2014) |
| MXAN_2437 | LWT | ECF Sigma Factors | This study |
| MXAN_2500 | Early aggregation | ECF Sigma Factors | This study |
| MXAN_2501 | Aggregate-reaggregate | NtrC-Like Activators | This study |
| MXAN_2516 | Immature aggregates | NtrC-Like Activators | This study |
| MXAN_2654 | Early aggregation | ABC Transporter | (Yan *et al,* 2014) |
| MXAN_2711 | Variable | One Component | (Ritchie *et al,* 2021) |
| MXAN_2783 | Early aggregation | ABC Transporter | (Yan *et al,* 2014) |
| MXAN_2794 | Immature aggregates | One Component | This study |
| MXAN_2795 | LWT | ABC Transporter | (Yan *et al,* 2014) |
| MXAN_2831 | LWT | ABC Transporter | (Yan *et al,* 2014) |
| MXAN_2832 | LWT | ABC Transporter | (Yan *et al,* 2014) |
| MXAN_2833 | LWT | ABC Transporter | (Yan *et al,* 2014) |
| MXAN_2853 | Early aggregation | ABC Transporter | (Yan *et al,* 2014) |
| MXAN_2896 | LWT | One Component | This study |
| MXAN_2929 | LWT | ECF Sigma Factors | This study |
| MXAN_2949 | Aggregate-reaggregate | ABC Transporter | (Yan *et al,* 2014) |
| MXAN_2951 | Early aggregation | ABC Transporter | (Yan *et al,* 2014) |
| MXAN_3095 | Early aggregation | NtrC-Like Activators | This study |
| MXAN_3142 | Immature aggregates | One Component | This study |
| MXAN_3151 | Immature aggregates | One Component | This study |
| MXAN_3208 | Immature aggregates | ABC Transporter | (Yan *et al,* 2014) |
| MXAN_3209 | Early aggregation | ABC Transporter | (Yan *et al,* 2014) |
| MXAN_3214 | Fall apart | NtrC-Like Activators | (Ritchie *et al,* 2021) |
| MXAN_3240 | LWT | One Component | This study |
| MXAN_3256 | No aggregation | ABC Transporter | (Yan *et al,* 2014) |
| MXAN_3257 | No aggregation | ABC Transporter | (Yan *et al,* 2014) |
| MXAN_3258 | No aggregation | ABC Transporter | (Yan *et al,* 2014) |
| MXAN_3333 | Variable | NtrC-Like Activators | This study |
| MXAN_3339 | LWT | ABC Transporter | (Yan *et al,* 2014) |
| MXAN_3381 | Aggregate-reaggregate | NtrC-Like Activators | This study |
| MXAN_3418 | LWT | NtrC-Like Activators | This study |
| MXAN_3426 | Early aggregation | ECF Sigma Factors | This study |
| MXAN_3429 | Early aggregation | One Component | This study |
| MXAN_3443 | Immature aggregates | One Component | This study |
| MXAN_3648 | Variable | ABC Transporter | (Yan *et al,* 2014) |
| MXAN_3650 | LWT | ABC Transporter | (Yan *et al,* 2014) |
| MXAN_3686 | Early aggregation | ECF Sigma Factors | This study |
| MXAN_3692 | No aggregation | NtrC-Like Activators | This study |

| | | | |
|---|---|---|---|
| MXAN_3702 | No aggregation | One Component | (Ritchie *et al,* 2021) |
| MXAN_3711 | Immature aggregates | One Component | This study |
| MXAN_3717 | No aggregation | ABC Transporter | (Yan *et al*, 2014) |
| MXAN_3718 | Fall apart | ABC Transporter | (Yan *et al*, 2014) |
| MXAN_3773 | LWT | ABC Transporter | (Yan *et al*, 2014) |
| MXAN_3811 | LWT | NtrC-Like Activators | This study |
| MXAN_3908 | LWT | ABC Transporter | (Yan *et al*, 2014) |
| MXAN_3909 | Early aggregation | ABC Transporter | (Yan *et al*, 2014) |
| MXAN_3959 | Early aggregation | ECF Sigma Factors | This study |
| MXAN_3986 | LWT | ABC Transporter | (Yan *et al*, 2014) |
| MXAN_4042 | Immature aggregates | NtrC-Like Activators | This study |
| MXAN_4060 | LWT | One Component | This study |
| MXAN_4072 | Late aggregation | One Component | This study |
| MXAN_4110 | Late aggregation | One Component | This study |
| MXAN_4173 | LWT | ABC Transporter | (Yan *et al*, 2014) |
| MXAN_4196 | No aggregation | NtrC-Like Activators | (Ritchie *et al,* 2021) |
| MXAN_4199 | LWT | ABC Transporter | (Yan *et al*, 2014) |
| MXAN_4240 | LWT | NtrC-Like Activators | This study |
| MXAN_4247 | Late aggregation | One Component | This study |
| MXAN_4252 | Late aggregation | NtrC-Like Activators | This study |
| MXAN_4261 | LWT | NtrC-Like Activators | This study |
| MXAN_4263 | LWT | One Component | This study |
| MXAN_4309 | Early aggregation | ECF Sigma Factors | This study |
| MXAN_4316 | Early aggregation | ECF Sigma Factors | This study |
| MXAN_4339 | LWT | NtrC-Like Activators | This study |
| MXAN_4356 | LWT | One Component | This study |
| MXAN_4471 | LWT | One Component | This study |
| MXAN_4523 | LWT | ABC Transporter | (Yan *et al*, 2014) |
| MXAN_4580 | Early aggregation | NtrC-Like Activators | This study |
| MXAN_4622 | LWT | ABC Transporter | (Yan *et al*, 2014) |
| MXAN_4662 | Immature aggregates | ECF Sigma Factors | This study |
| MXAN_4665 | LWT | ABC Transporter | (Yan *et al*, 2014) |
| MXAN_4716 | Fall apart | ABC Transporter | (Yan *et al*, 2014) |
| MXAN_4733 | Early aggregation | ECF Sigma Factors | This study |
| MXAN_4749 | LWT | ABC Transporter | (Yan *et al*, 2014) |
| MXAN_4750 | LWT | ABC Transporter | (Yan *et al*, 2014) |
| MXAN_4785 | Late aggregation | NtrC-Like Activators | This study |
| MXAN_4790 | Variable | ABC Transporter | (Yan *et al*, 2014) |
| MXAN_4899 | Fall apart | NtrC-Like Activators | This study |
| MXAN_4949 | Early aggregation | ECF Sigma Factors | This study |
| MXAN_4977 | Early aggregation | NtrC-Like Activators | This study |

| | | | |
|---|---|---|---|
| MXAN_4983 | Variable | NtrC-Like Activators | This study |
| MXAN_4987 | LWT | ECF Sigma Factors | This study |
| MXAN_5029 | No aggregation | One Component | This study |
| MXAN_5041 | Immature aggregates | NtrC-Like Activators | This study |
| MXAN_5048 | Late aggregation | NtrC-Like Activators | This study |
| MXAN_5101 | Early aggregation | ECF Sigma Factors | (Ritchie *et al,* 2021) |
| MXAN_5124 | Fall apart | NtrC-Like Activators | (Ritchie *et al,* 2021) |
| MXAN_5128 | LWT | One Component | This study |
| MXAN_5153 | Early aggregation | NtrC-Like Activators | (Ritchie *et al,* 2021) |
| MXAN_5245 | Late aggregation | ECF Sigma Factors | This study |
| MXAN_5263 | No aggregation | ECF Sigma Factors | (Ritchie *et al,* 2021) |
| MXAN_5271 | No aggregation | One Component | This study |
| MXAN_5276 | LWT | ABC Transporter | (Yan *et al,* 2014) |
| MXAN_5305 | LWT | One Component | This study |
| MXAN_5356 | Early aggregation | One Component | This study |
| MXAN_5379 | LWT | ABC Transporter | (Yan *et al,* 2014) |
| MXAN_5410 | Early aggregation | ECF Sigma Factors | (Ritchie *et al,* 2021) |
| MXAN_5480 | Early aggregation | One Component | This study |
| MXAN_5492 | LWT | One Component | This study |
| MXAN_5503 | LWT | ABC Transporter | (Yan *et al,* 2014) |
| MXAN_5506 | Early aggregation | ECF Sigma Factors | This study |
| MXAN_5545 | Variable | One Component | This study |
| MXAN_5547 | LWT | One Component | This study |
| MXAN_5584 | LWT | ABC Transporter | (Yan *et al,* 2014) |
| MXAN_5680 | Variable | NtrC-Like Activators | (Ritchie *et al,* 2021) |
| MXAN_5731 | Immature aggregates | ECF Sigma Factors | This study |
| MXAN_5777 | Variable | NtrC-Like Activators | (Ritchie *et al,* 2021) |
| MXAN_5853 | Variable | NtrC-Like Activators | This study |
| MXAN_5879 | Fall apart | NtrC-Like Activators | (Ritchie *et al,* 2021) |
| MXAN_5894 | Variable | One Component | (Ritchie *et al,* 2021) |
| MXAN_6000 | Late aggregation | ABC Transporter | (Yan *et al,* 2014) |
| MXAN_6058 | Variable | ECF Sigma Factors | This study |
| MXAN_6149 | LWT | One Component | This study |
| MXAN_6157 | LWT | One Component | This study |
| MXAN_6161 | LWT | One Component | This study |
| MXAN_6167 | Variable | One Component | This study |
| MXAN_6173 | Early aggregation | ECF Sigma Factors | (Ritchie *et al,* 2021) |
| MXAN_6206 | Variable | One Component | This study |
| MXAN_6251 | Variable | One Component | This study |
| MXAN_6402 | LWT | ABC Transporter | (Yan *et al,* 2014) |
| MXAN_6426 | No aggregation | NtrC-Like Activators | (Ritchie *et al,* 2021) |

| | | | |
|---|---|---|---|
| MXAN_6461 | Fall apart | ECF Sigma Factors | This study |
| MXAN_6468 | Variable | One Component | This study |
| MXAN_6475 | Early aggregation | ABC Transporter | (Yan *et al*, 2014) |
| MXAN_6479 | LWT | One Component | This study |
| MXAN_6486 | LWT | One Component | This study |
| MXAN_6518 | Variable | ABC Transporter | (Yan *et al*, 2014) |
| MXAN_6549 | Late aggregation | One Component | This study |
| MXAN_6551 | LWT | ABC Transporter | (Yan *et al*, 2014) |
| MXAN_6646 | LWT | One Component | This study |
| MXAN_6653 | No aggregation | One Component | This study |
| MXAN_6759 | Immature aggregates | ECF Sigma Factors | This study |
| MXAN_6833 | Variable | One Component | This study |
| MXAN_6889 | No aggregation | One Component | (Ritchie *et al*, 2021) |
| MXAN_6967 | Late aggregation | One Component | This study |
| MXAN_7072 | Variable | One Component | This study |
| MXAN_7078 | LWT | One Component | This study |
| MXAN_7214 | Early aggregation | ECF Sigma Factors | This study |
| MXAN_7289 | Immature aggregates | ECF Sigma Factors | This study |
| MXAN_7312 | LWT | One Component | This study |
| MXAN_7316 | LWT | One Component | This study |
| MXAN_7322 | Late aggregation | One Component | This study |
| MXAN_7326 | LWT | ECF Sigma Factors | This study |
| MXAN_7440 | No aggregation | NtrC-Like Activators | (Ritchie *et al*, 2021) |
| MXAN_7454 | Early aggregation | ECF Sigma Factors | This study |

### *References*


Bradski G (2000) The OpenCV Library. *Dr Dobb's J Softw Tools*

Madeira F, Pearce M, Tivey ARN, Basutkar P, Lee J, Edbali O, Madhusoodanan N, Kolesnikov A & Lopez R (2022) Search and sequence analysis tools services from EMBL-EBI in 2022. *Nucleic Acids Res*: 1–4

Ritchie LJ, Curtis ER, Murphy KA & Welch RD (2021) Profiling Myxococcus xanthus Swarming Phenotypes through Mutation and Environmental Variation. *J Bacteriol* 203

Yan J, Bradley MD, Friedman J & Welch RD (2014) Phenotypic profiling of ABC transporter coding genes in Myxococcus xanthus. *Front Microbiol* 5: 1–12
